# Modular protein-oligonucleotide signal exchange

**DOI:** 10.1101/825422

**Authors:** Deepak K. Agrawal

**Affiliations:** Department of Chemical and Biomolecular Engineering, Johns Hopkins University, 3400 N Charles St, Baltimore, Maryland 21218, USA

## Abstract

The ability to detect a protein selectively and produce a predicted signal in real time is a long-lasting engineering challenge in the field of biochemistry. Such a mechanism typically requires a sensing module to recognize the input protein and a translation module to produce a programmable output signal that reflects the concentration of the input. Here we present a generic biomolecular reaction process that exchanges the concentration of an input protein with a DNA oligonucleotide. This approach uses the unique characteristic of DNA oligonucleotide aptamer that can either bind to a specific protein or to a complementary DNA oligonucleotide reversibly. We then pass the information of the protein concentration to the output signal through DNA strand displacement reactions. Using this strategy, we design and characterize four different exchange processes that can produce modular DNA oligonucleotides in response to different proteins such as clinically important human α-thrombin and vascular endothelial growth factor (VEGF). These exchange processes are capable of real time sensing and are modular such that they can be used for concurrent detection of different proteins with well-defined input-output characteristics. The novelty and simplicity of our approach encourage to develop advanced biochemical systems for point-of-care testing of infectious diseases and treatments.

## INTRODUCTION

Cell continuously responds to its environment, mainly by producing and degrading a variety of protein molecules that play a central role in metabolic reactions. These reactions are often performed by transcriptional networks, determining the rate at which specific proteins are produced and then used downstream as transcriptional factors to control the production of other proteins. This results in complex pathways *in vivo* wherein a protein may control many downstream pathways or many proteins controlling a single pathway (1-3). Such a complex operation *in vitro* may only be possible using a modular approach that could detect a specific protein and in response, capable of producing an output molecule that could easily be programmed.

Interestingly, there is a class of short single stranded oligonucleotides that bind to target molecules with high affinity and specificity. These DNA and RNA oligonucleotides are known as aptamers and often fold into secondary or tertiary structures upon binding to a specific target (4-6). Because a DNA oligonucleotide can easily be realized and used in lab compared to RNA, various DNA aptamer based biosensing assays have been developed (7-12). Moreover, due to the predictable nature of the DNA molecule, governed by the Watson-Crick base-pairing, short DNA oligonucleotides have emerged as a powerful programmable material to develop various biochemical circuits capable of mimicking digital and analog circuit operations found in electronic circuits (13, 14), including a programmable DNA circuit capable of controlling a protein functionality (15). However, none of these approaches resulted in a generalized modular scheme that could be readily available to translate the concentration of the input protein into a programmable molecule with a well-defined response.

In a cell, different signal pathways are used to respond to various external and internal activities. Because these signal pathways can work together without effecting the operation of other, cell can perform complex processes such as growth, metabolism and cell division effectively (16). A mechanism that can perform similar operations *in vitro*, requires a modular reaction network to exchange the concentration of different proteins into one or different to a DNA output strand modularly and then, the output of this reaction network may be programmed using downstream strand displacement reactions, which are based on hybridization and displacement of short single strands, to control other molecules.

In this paper, we construct a simple generic platform that modularly produces a DNA oligonucleotide in exchange for an input protein. Our strategy relays in series of toehold exchange strand displacement reactions that provide enough freedom to select the sequence of the output DNA oligonucleotide independent of the sequence of DNA aptamer. This unique characteristic of the exchange process allows to detect different proteins using their DNA aptamers and can produce output DNA oligonucleotide of virtually any sequence. We exploit the natural interaction of a DNA aptamer with a specific protein to infer the concentration of the protein and then propagate this information to the output using reversible strand displacement cascade for real time detection. The modular design of the reaction network enables us to produce the same DNA output strand in the presence of two different proteins, and also different DNA output strands in response to the same protein concurrently.

## DESIGN

Our goal was to construct a reversible process of biomolecular reactions where the amount of a specific DNA oligonucleotide is dynamically modulated by an input protein. The process output then may be used as the input signal to downstream chemical logic (17), amplification (18) or other control processes (19).

Here, we present a modular exchange process that meets these requirements for a variety of protein inputs and virtually arbitrary short oligonucleotide sequence outputs. The requirements for developing such a system for a specific protein or other input molecules is the existence of a DNA aptamer that reversibly binds to this target. A reversible binding with a well-known relation between the aptamer and its target is required for real time sensing. This can allow obtaining a well-defined predictable change in the aptamer concentration in response to the input molecule. We can then incorporate a downstream process for producing a common output signal in response to different proteins (Figure 1A), and also different output signal for the same protein (Figure 1B). Such a reaction network, in principle, could then be used to control multiple downstream processes in conjunction, such as by serving as an input to strand displacement processes (20, 21). Further, these networks are modular, such that reaction systems that translate the concentrations of multiple proteins into the concentrations of numerous oligonucleotides can operate in the same test tube.

**Figure 1.**
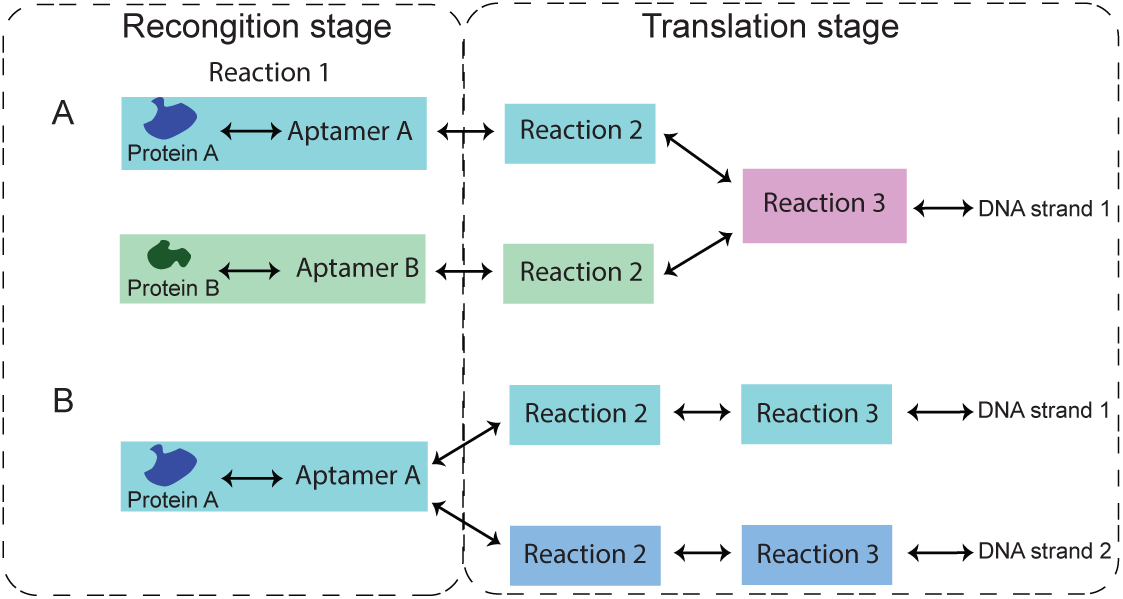
Block diagram representation of protein-oligonucleotide signal exchange processes. Each exchange process consists of a recognition stage where aptamer targets a specific protein and a translation stage where reversible toehold-mediated strand-displacement reactions process the free aptamer concertation into an output DNA strand quickly. Here, input to Reaction 2 is the free aptamer strand; the presence of the protein reduces the concentrations of free aptamer, which in turn changes the concentration of the output DNA strand. (a) Different protein inputs can modulate the concentration of the same DNA strand output using two exchange processes with different inputs and the same output. Concurrent operation of these exchange processes requires that the free aptamers of each pathway can interact only with the respective downstream Reaction 2. (b) Multiple exchange processes can produce two oligonucleotide outputs with no sequence homology in response to the same input protein. Such an operation requires that the reactions involved in the translation stage to exhibit minimal crosstalk.

As a starting point, we chose to divide the exchange process into two stages based on their operations. The first stage is a recognition stage that uses reversible aptamer-target binding (Figure 2, pink) and the second stage is a translation, where a change in the concentration of unbound aptamer is processed into a change in the free concentration of a DNA output strand that has no sequence in common with the aptamer sequence. This operation relies on the high affinity binding of the DNA aptamer to the input protein such that protein sequesters the aptamer quickly before it can react with any other species in the same mixture. Assuming this reaction is close to equilibrium, the amount of free aptamer is a well-defined function of the initial aptamer and protein concentration at the aptamer’s equilibrium dissociation constant (K_d_). Using the K_d_ value, one can choose the aptamer concentration such that small changes in the concentration of the input can produce measurable changes in the aptamer concentration.

**Figure 2.**
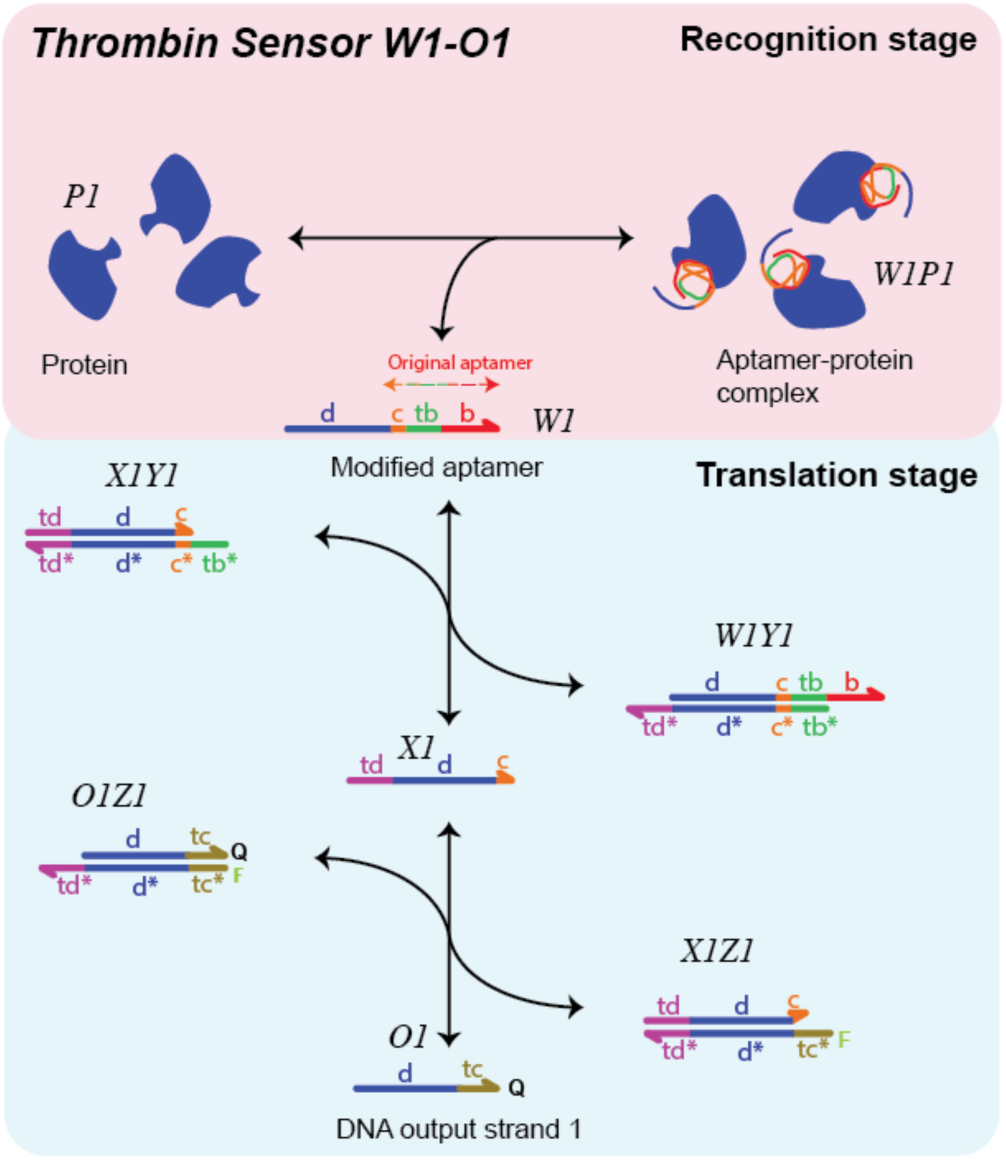
Schematic of a protein-oligonucleotide signal exchange process for dynamically reporting protein concentration as the concentration of an output oligonucleotide with the designed sequence. The concentration of the output strand O1 is modulated by the concentration of the input protein P1. The folded aptamer (W1) binds to the input protein (thrombin for *Sensor W1-O1*) to form an aptamer-protein (W1P1) complex. Aptamer strands that are not bound to the protein can react with the X1Y1 complex, shifting the equilibrium of a reversible reaction so as to change the concentration of free O1. [O1]_free_ ranges from [O1]_total_ (when [P1]=0) to 0 (when [P1] -> ∞) that is determined by free protein concentration. O1 has no sequence domains in common with the aptamer (SI Note S2), so it can be chosen largely freely in order to couple it to a previously designed downstream process. The forward and reverse reactions within the strand displacement cascade are each relatively fast because they are mediated by 4-5 base pair tb, or 5 base pair tc or 6 base pair (bp) toehold binding [22]. Typically, for reversible toehold-mediated strand-displacement reactions, these rates vary from 1 M^-1^s^-1^ (zero toehold) to 6 ×10^6^ M^-1^s^-1^ (7 toehold or longer) (22). For *Thrombin Sensor W1-O1*, domains d, c, b and tb are, respectively, 15, 2, 8 and 5 bp long. Sequences for all the exchange process are listed in SI Table S3, S6, S7 and S8. Here, [] represents the molar concentration of the species.

Instead of using the free concertation of the DNA aptamer as an output signal, we incorporate a translation stage that produces a DNA output strand that reflects the free concentration of aptamer (Figure 2, cyan). The idea is that the concentration of this free output strand should vary precisely with the precise concentration of the input protein, without any requirement that the output strand shares a sequence in common with the aptamer. This is required to develop a modular exchange process that could be readily used to detect different proteins. To achieve this functionality between input and output, we use a reversible, toehold-mediated strand displacement cascade that processes the free concentration of any desired output strand such that it reflects the concentration of free aptamer.

### Designing the exchange process

To design the recognition stage that can process a protein concertation into the free aptamer concertation, we start by assigning the aptamer core sequence into c, b and tb domains. Domains c and b bind to the respective complementary domains irreversibly while tb is a short toehold domain that serves as a nucleation site for a toehold-mediated branch migration reaction with the components of the translation portion of the process (22). To perform sequence translation, we added a new domain, the d domain, at the 5’-end of the original aptamer. The d domain mediates the interaction of the free aptamer with the complexes that perform the sequence translation and is part of the sequence of the output strand in the 2-stage translation reaction that we designed. An interaction between the domain d and the core aptamer sequence could lead to an undesirable folding and/or reducing the affinity of the aptamer towards its target (9). To avoid this, domain d was composed solely of A’s and T’s and the secondary structure modelling using NUPACK (23) predicted no interaction between the d domain and its respective core aptamer sequence for each modified aptamer.

The translation stage requires to provide sequence independence between the aptamer sequence and the sequence of the output strand. We achieve this by a two-step strand-displacement (Figure 2, cyan). In the first step, the aptamer binds to a complex X1Y1 by its toehold domain tb and releases strand X1. Strand X1 contains domain d as well as domain c which is part of the core aptamer sequence, so it has some sequence overlap with the aptamer. Strand X1 can then react with the O1Z1 complex, freeing the output strand O1 and producing X1Z1 complex where the c domain of the aptamer is single-stranded. O1 has no overlap sequence with the core aptamer sequence. This could allow producing an output signal in response to different input aptamer that has the same d domain (Figure 1A), and different output signals in response to an input aptamer that has different d domains (Figure 1B).

A reliable dynamic sensing requires all the reactions in the exchange network to be reversible and fast so that the output of the exchange network can converge quickly to an equilibrium in response to any change in the input concentration such that the initial and final concentrations of the output strand correspond to the input protein concentration via a well-defined dose-response curve. Therefore, using toehold-mediated strand-displacement process, we design reversible strand-displacement reactions wherein rates of these reactions were controlled by appropriately choosing the length of toehold domains. Typically, for reversible toehold-mediated strand-displacement reactions, these rates vary from 1 M^-1^s^-1^ (zero toehold) to 6 ×10^6^ M^-1^s^-1^ (7 toehold or longer) (22).

### Kinetic modelling of the exchange process

Our goal in developing modular protein-oligonucleotide signal exchange process was not to create a mechanism that can not only detect the presence or absence of an input protein, but also to precisely modulate the concentration of the output oligonucleotide signal in response to variations in the input protein concentration. Our goal, thus, was to produce a process with a well-defined dose-response curve. Such characteristic could allow the exchange process to translate input concentrations of many kinds of molecules into oligonucleotide concentrations, where these oligonucleotides could serve as inputs for a growing variety of analytical systems for molecular logic (24), signal modulation (25), or within synthetic gene networks (26) where the concentration of the oligonucleotide is important for determining the result of the process.

To establish the precise relationship between process input and output, we developed a modelling framework that relied on known equilibrium constants and/or reaction rate constants of the reactions involved in the exchange process. These parameters were then used to model of mass-action kinetics of coupled chemical reactions:

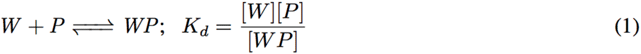

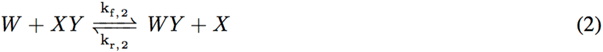

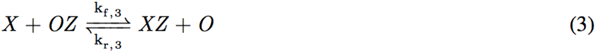

Here and elsewhere W, P, WP, XY, WY, X, OZ, XZ and O species are referred in general for any exchange network, and [] represents the molar concentration of the species. k_f_ and k_r_ represent forward and reverse rate constants respectively. Reaction 1 corresponds to the recognition stage of the exchange process where DNA aptamer (W) binds to a protein (P) with typically a smaller value of dissociation constant (K_d_), suggesting a high affinity between aptamer and protein. This means that the final change in the concentration of the aptamer from its initial value once the input protein was added can be related accurately using the K_d_ value. To model this reaction for each exchange process, we relied on the literature for K_d_ value and kinetic rates.

Reactions 2 and 3 correspond to the translation stage that uses a strand displacement technique to reflect the changes of free aptamer concentration into the concentration of the output DNA strand (O). To be able to model these changes accurately, we calculated the rates associated with these reactions using the kinetic model of the strand displacement reactions that are mediated by the interaction of the toeholds (22). This model requires binding energies for each complex (XY, WY, OZ and XZ) involved in the reactions as the input. To calculate these binding energies, we used NUPACK simulation package (23) and then used the kinetic model to calculated the rates (k_f,2_, k_r,2_, k_f,3_, k_r,3_) of Reactions 2 and 3 for each exchange process (see SI Note S1). The exchange process we designed uses sequential reactions wherein the output of one reaction serves as input to another reaction. Therefore, simple equilibrium analysis may not be accurate to predict the output signal with changes in the input aptamer or protein concentrations. Therefore, we used a dynamic ordinary differential equations (ODEs) model that uses forward and reverse rates of each reaction to model the entire exchange network (see SI Note S1).

## RESULTS

### Designing and characterizing Thrombin Sensor W1-O1

To test whether the exchange process could be used to translate the concentration of an input protein into an output DNA strand that shares no sequence with the original sequence of the aptamer, we first designed *Thrombin Sensor W1-O1*. This process targets human α-thrombin, a serine protease that is involved in activating coagulation cascade (28). To detect thrombin, we adapted the previously studied DNA 15-mer TBA15 as a thrombin binding aptamer (29) that controls the concentration of a DNA output strand (O1) through translation stage. Following our design methods (described above), we started with assigning the specific sequence of the 15-mer TBA15 aptamer into c, tb and b domains and added a designed 15 base pair d domain at its 5’-end that was contained in the output strand. We then designed X1Y1 and O1Z1 complexes such that only the modified aptamer (W1) interacts with the thrombin, but the free W1 can still interact with X1Y1 (see SI Note S2). We start by characterizing the recognition and translation stages separately and then test the response of the exchange process as a whole.

For each of the reactions in the exchange process to function without undesired crosstalk. In particular, if the single-stranded b domain of the aptamer, presented by the W1Y1 complex, or the single-stranded c domain of the aptamer, presented by the X1 strand could react with thrombin, the free aptamer concentration would not be solely determined by the K_d_ of the aptamer, but also by the reactions between the thrombin and one or more of these complexes. These interactions may be avoided by choosing b and c domains such that these domains do not fold into tertiary structures (like G-quadruplex) that may have some affinity towards thrombin. Therefore, we considered structural data for TBA15 (27) in deciding how to divide the domains and chose them so that b and c domains could not form the G-quadruplex required for thrombin affinity. We also selected domain divisions so that no one domain was too long: b is 8 nucleotides, tb is 5 nucleotides and c is 2 nucleotides (SI Note S2).

### Binding of the modified aptamer to the input protein in the recognition stage

Our first step in characterizing the recognition stage of *Thrombin Sensor W1-O1* was to verify that the only the modified aptamer that contains the TBA15 sequence and the d domain, still bound with the high affinity to thrombin. To test this, we used a non-denaturing electrophoresis mobility shift assay (EMSA) that measures the ability of thrombin to enter the gel only when thrombin bound to the aptamer (27). We added different concentrations of thrombin (1, 2.5 and 5 µM) or control with 5 µM of 15-mer TBA15 (see Methods). Similar mixes were made for modified TBA15 (W1). In both the cases, we found that the fraction of the aptamer (either TBA15 or W1) that was bound to thrombin increased as the thrombin concentration increased, resulting in a darker band (SI Fig. S1).

Moreover, a higher molar ratio between the aptamer and thrombin resulted in a better migration of these bands, indicating the higher amount of the aptamer bound to thrombin in the gel matrix. For both the aptamers, we observed almost a similar gel matrix response, suggesting that the K_d_ of the modified TBA15 for thrombin should be close to the K_d_ of the original 15-mer TBA15. We then tested whether other species in the exchange process such as W1Y1 complex through b doming and X1 strand through c domain could interact with thrombin. Such interactions can lead to a scenario where the free aptamer concentration would not be solely determined by the K_d_ of the aptamer, but also by the reactions between the thrombin and one or more of these complexes. No thrombin was detected in the presence of W1Y1 complex or X1 strand (SI Fig. S2), suggesting that b or c domain alone has no affinity towards the thrombin.

### Characterizing translation stage response in protein free environment

Our goal was to construct an exchange process where the concentration of the output strand is a known, reproducible function of the input protein. We have already qualitatively established that the modified aptamer can still bind strongly with thrombin and no other complexes can interact with thrombin except the aptamer. Our next goal was to characterize the response of the translation portion of the exchange process, where the free aptamer interacts with downstream complexes to modulate the concentration of free output strand.

To test what concentrations of free aptamer produce what concentrations of the output strand, we varied the concentration of W1 while having the initial concentrations of X1Y1 and O1Z1 complexes to be 100 nM each. To be able to read out the concentration of the output strand (O1) in real time, we attached a fluorescent molecule to the 5’ end of the Z1 strand (FAM unless otherwise mentioned) and a quencher to the 3’ end of the O1 strand, which is hybridized to Z1 when it is not free (see Methods). The fluorescence of the mixture therefore reflects the concentration of free O1 (Figure 3A, see SI Note S3).

**Figure 3.**
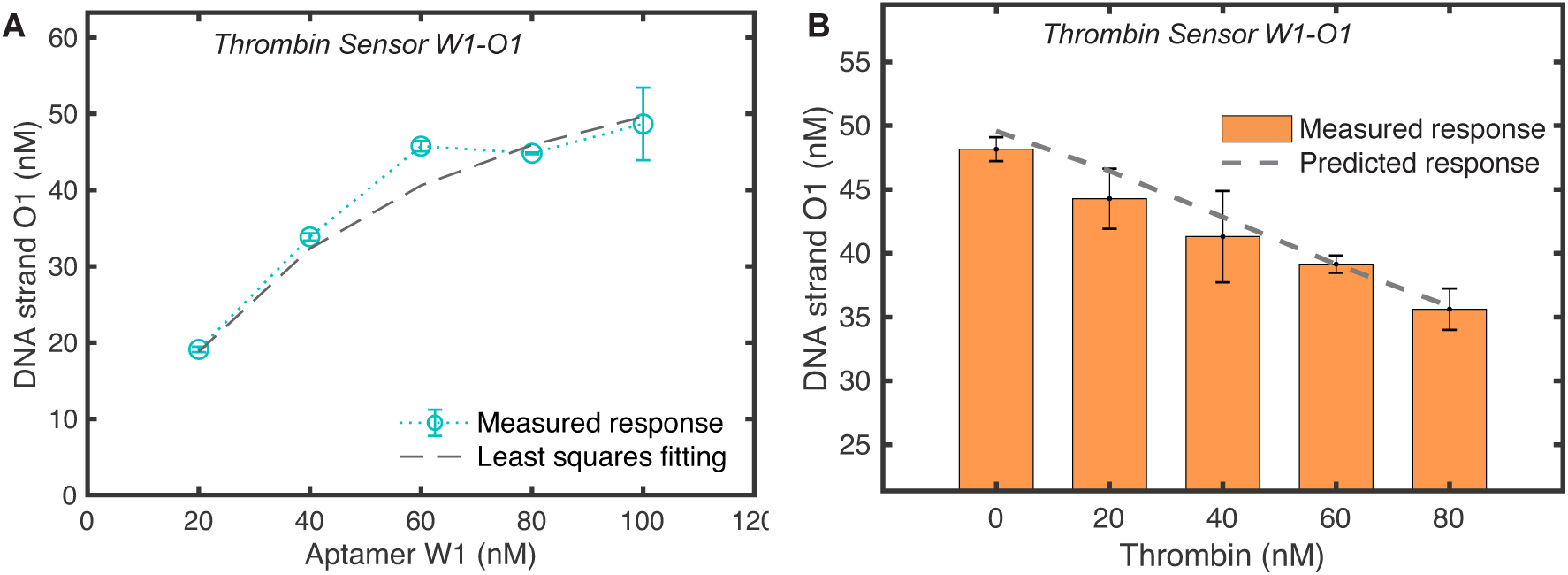
Input protein concentration controls output oligonucleotide concentration through the free aptamer concentration. **(A)** The equilibrium concentration of O1 in a protein free environment for *Thrombin Sensor W1-O1* at different concentrations of W1 and [X1Y1]=[O1Z1]=100 nM. **(B)** The equilibrium concentration of O1 to different concentrations of thrombin where [W1]=[X1Y1]=[O1Z1]=100 nM. For both experiments (and elsewhere in the paper), a calibration process using known concentrations of O1 was used to convert the measured fluorescence intensities into O1 concentrations (see SI Note S3 for details). The error bars here and elsewhere were determined using the standard error of the mean of three or more measured responses. Here and elsewhere, a dynamic ODE model was used to determine the response of the exchange process without and with protein. The reaction rates were determined using the kinetic model of toehold mediated strand-displacement (22) except the forward rate constant of Reaction 2 that was determined using the least squares fitting of the measured protein free response. The W1-P1 interaction was modelled using the K_d_ value and forward rate constants reported in (30, 31) (see SI Note S1).

To predict the concentration of O1 in response to different concentration of W1, we first calculated the rate constants of Reactions 2 and 3 for *Thrombin Sensor W1-O1* using the kinetic model of strand displacement reaction that described earlier (see Design section). We then fed these values in an ODE model for the exchange process (see SI Note S1) to determine O1 concentration with changes W1 concentration. Our modelling predicted a higher amount of O1 than were measured (SI Fig. S6). We hypothesized that this discrepancy might be caused by the G-quadruplex structure within W1, preventing interaction between W1 and the X1Y1 through hybridization of the tb domain, which initiates the branch migration of domain c. This hindrance would be expected to reduce the forward rate constant (k_f,2_) of Reaction 2; furthermore, this reduced rate would not be predicted by NUPACK, which does not consider the energetics of tertiary structures such as the G-quadruplex.

To test our hypothesis, we directly measured the changes in the concentration of X1 strand when different concentrations of W1 were added to mixtures containing 250 nM of X1Y1 complex that has a fluorophore molecule to the 5’ end of X1 strand and a quencher to the 3’ end of Y1 strand (see Methods, SI Fig. S7A). We found that for a k_f,2_ value of 5 × 10^4^ M^-1^s^-1^, the measured response of Reaction 2 alone closely follows the model data (SI Fig. S7B). This value is much smaller than the rate that was calculated using the strand displacement model (2.83 × 10^6^ M^-1^s^-1^). We then used the least squares fitting method to find a k_f,2_ value that would allow a best fit between the model data and the measure protein free response and found that this revised model showed close agreement with the experimental data for a k_f,2_ value of 6.4 × 10^4^ M^-1^s^-1^ (Figure 3A), which is close to the one we measured earlier.

### Detecting different protein input concentrations

Having validated that the operation of recognition and translation stages of *Thrombin Sensor W1-O1*, our next goal was to characterize the behaviour of the full exchange process. In this process, an incremental change in the concentration of thrombin should incrementally reduce the amount of O1. These changes should be predicted by a model that incorporates the rates of the reaction involved in the translation stage and the rates of TBA15-thrombin interaction that borrowed from literature (30, 31) (see SI Table S2). To test this, we first incubated 100 nM of W1 with different amounts of thrombin (ranging from 0-80 nM) for 30 min and then added this mixture to a solution containing 100 nM of each X1Y1 and O1Z1 and measured the steady state fluorescence intensities for each thrombin concentration mixture. Converting these intensities into [O1] values, (see SI Note S3), we observed that [O1] at steady state was almost identical to the predicted data through our model for each of the thrombin concentrations tested (Figure 3B). Multiple tests of the exchange process produced very similar input-output responses.

The exchange process should be able to reliably produce the predictable changes in O1’s concentration when the thrombin concentration was added in smaller or in larger quantities compared to the values reported in Figure 3B. To verify this, we tested the response of *Thrombin Sensor W1-O1* at different ranges of input thrombin concentrations. During these tests, to modulate the changes in the concentration of O1 strand accurately, we used different initial concentrations of the sensor components (W1, X1Y1 and O1Z1) (see Methods). For each variation in thrombin concentration, a well-predicted response was observed (SI Fig. S8), thus validating not just our modelling approach but also the fact that we can accurately predict the input-output response of this exchange process.

One of the main design considerations of the reaction cascade is that the free aptamer concentration should entirely control the steady state output strand’s concentration. This requires that thrombin should not interact with any other component of the translation system. For that, thrombin should not interact with any other component of the translation system. We, therefore, tested the operation of the sensor in the absence of aptamer and observed a negligible change in the output as a function of thrombin concentrations (SI Fig. S8A). Because the thrombin stock used was dissolved in 50% H_2_O/Glycerol mix, we also tested whether such a mixture alone would shift fluorescence values, but we observed no significate change [O1] (SI Fig. S9B).

### Detecting dynamic changes in thrombin concentration overtime

The reversible nature of the exchange process should allow the concentration of the output strand to change in response to changes in protein concentration by changing the output strand’s equilibrium concentration. To test whether such updates occur, we first characterized how *Thrombin Sensor W1-O1* responded to increases in thrombin concentration in the same test tube overtime. For this, we varied the thrombin concentration in higher amounts than the standard sensing assay so that each change in the fluorescence signal could be tracked accurately. We first prepared a mixture consisting of 500 nM W1, X1Y1 and O1Z1 each. We then added thrombin to a final concentration of 100 nM to the mixture and measured the fluorescence at steady state (see Methods). We then added more thrombin so that the final concentration of thrombin in the mixture had increased to 200 nM. The output fluorescence decreased after thrombin was added, and each time after more thrombin was added in steps up to a final concentration of 400 nM (Figure 4A).

**Figure 4.**
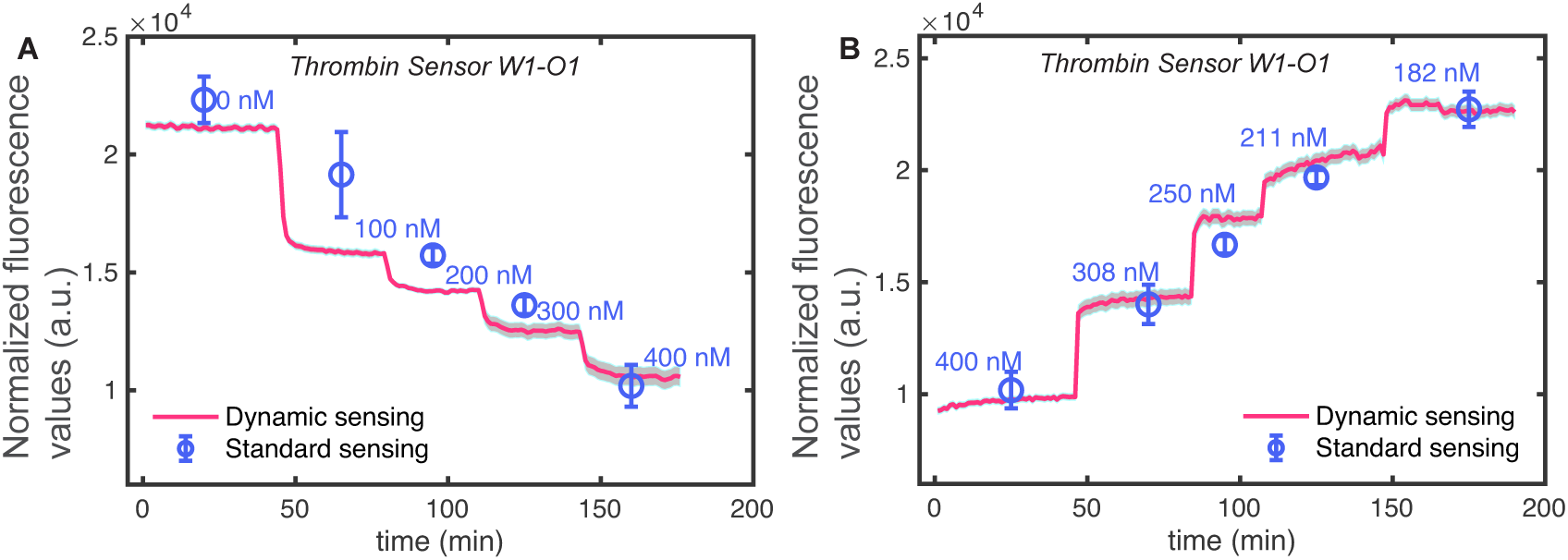
A signal exchange process can change its output value in response to changes in protein concentration in real time. Fluorescence response of *Thrombin Sensor W1-O1* as the function of thrombin concentration that was either increased or decreased overtime in a single mixture through the addition of thrombin or dilution of sensor components (W1, X1Y1 and O1Z1) respectively (dynamic sensing assay in Methods). These responses are compared with measurements where for each thrombin concertation a separate mixture was made (standard sensing assay in Methods) while maintaining the same volume and molarity of thrombin and sensor components as in the dynamic sensing case (see Methods). **(A)** To test how the reversible exchange process responds to increases in thrombin concentration, we mixed first X1Y1 and O1Z1 at 500 nM each and then added 500 nM of W1. We then added small volumes of thrombin from a concentrated stock in steps to increase thrombin concentration such that the sensor component’s concentration was not significantly affected because of the thrombin addition (see Methods). **(B)** To test how the exchange process responds to decreases in thrombin concentration, 500 nM of X1Y1 and O1Z1 were mixed and then added 500 nM of W1 followed by 400 nM of thrombin (first step). To reduce the thrombin concertation without affecting the concertation of sensor components, we then diluted this mixture by adding a fixed volume from a stock of sensor components containing [W1]=[X1Y1]=[O1Z1]=500 nM several times.

To determine whether the exchange process can also response to decreases in the thrombin concentration, we prepared a mixture consisting of 500 nM W1, X1Y1 and O1Z1 species each to which we initially added 400 nM of thrombin. To decrease the concentration of the thrombin without changing the effective concentration of the sensor components (W1, X1Y1 and O1Z1), we added a mixture containing 500 nM of the sensor components such that the final concentration of thrombin in the mixture was 308 nM while the sensor components remained at 500 nM (see Methods). The steady state fluorescence value increased after dilution. Further dilution steps using sensor network mixture that decreased the concentration of thrombin to 250 nM, 211 nM and then 108 nM each showed a significant increase in fluorescence value (Figure 4B).

Because our fluorescence measurements depended not only on the concentration of the unquenched fluorescent species but also on the volume of the mixture, it was not possible to use a single set of calibration measurement to determine the concentration of O1 after each of the dilution steps. To determine whether the changes in fluorescence after dilution indicated that the exchange process reached the expected equilibrium, for each of the measured values performed above, we prepared mixtures of the sensor components (at 500 nM each) and thrombin such that the final concentration was the same as in the above experiments, but for each thrombin concentration we had a separate mixture that was made by following the standard protein sensing assay (see Methods). The fluorescence measurements of these mixtures were consistent with those achieved through either the addition of more thrombin or more sensor components, indicating that the dynamic response of the system shifted the concentration of the output strand to its new equilibrium concentration, as designed (Figure 4).

### Producing identical DNA outputs using different aptamers or a different input protein

The modular design of the exchange process suggests that it should be straightforward to design an exchange process that produces an output strand of the same sequence as *Thrombin Sensor W1-O1* in response to either a different input protein, or that uses a totally different aptamer in the recognition stage. To demonstrate this capacity of the exchange process, we designed *Thrombin Sensor W2-O1*, that detects thrombin concentrations but does so using the DNA 29-mer DNA TBA29 to target thrombin, and *VEGF Sensor W3-O1* detects the concentration of vascular endothelial growth factor (VEGF) protein. Unlike TBA15 that binds to thrombin at its fibrinogen exosite and inhibits thrombin catalytic action to promote clotting, DNA aptamer TBA29 binds at the heparin exosite of thrombin with a K_d_ value of 500 pM (32) without affecting the thrombin catalytic activity (33). *VEGF Sensor W3-O1* uses a 25-mer 3R02 DNA aptamer that forms a stable G-quadruplex and binds to VEGF at its receptor-binding domain with a K_d_ value of 300 pM (34,35). The design process for these processes employed the design considerations followed when designing *Thrombin Sensor W1-O1* (see Design Section and SI Note S2 and S5). To produce the same DNA output strand (O1) from *Thrombin Sensor W2-O1* and *VEGF Sensor W3-O1* as in *Thrombin Sensor W1-O1*, the sequence of the d domain that was added in modified TBA29 and 3R02 aptamers was the same as was in modified TBA15 for *Thrombin Sensor W1-O1*.

To determine the relation between W2-O1 and W3-O1 for respectively *Thrombin Sensor W2-O1* and *VEGF Sensor W3-O1*, similar to *Thrombin Sensor W1-O1*, we initially characterized their responses at different W2 and W3 concentrations (SI Fig. S10A and S10B). To model these responses, we then calculated the reaction rates of the translation stage for each process (see SI Table S2) and found that using these rates, model data did not match with the measured response (SI Fig. S10A and S10B). Since both the aptamers used by these new processes have tertiary structures that might be affecting the rate of the strand displacement reaction between the respective aptamer and XY complex, we used the least squares fitting method to find the best fit for a k_f,2_ value at which the model data follows the protein free measured response of each sensor (SI Fig. S10C and S10D).

We then measured the response of *Thrombin Sensor W2-O1* and *VEGF Sensor W3-O1* separately to different concentrations of respective input proteins (Thrombin or VEGF) and in the presence of respective sensor components (W2, X2Y2, O1Z1 or W3, X3Y3, O1Z1) (Figure 5A and B respectively). For each exchange process, the measured O1 concentrations were then compared with the predicted response that used the same rates that were used to model the protein free response (SI Fig. S10C and S10D, SI Note S1). We found a close agreement between the modal and measured response for each exchange process (Figure 5A and B). By changing the concentrations of the sensor components, we were also able to modulate the concentration of O1 in response to a set of smaller and larger protein concentrations (SI Fig. S11 and S12) compared to Figure 5A and B.

**Figure 5.**
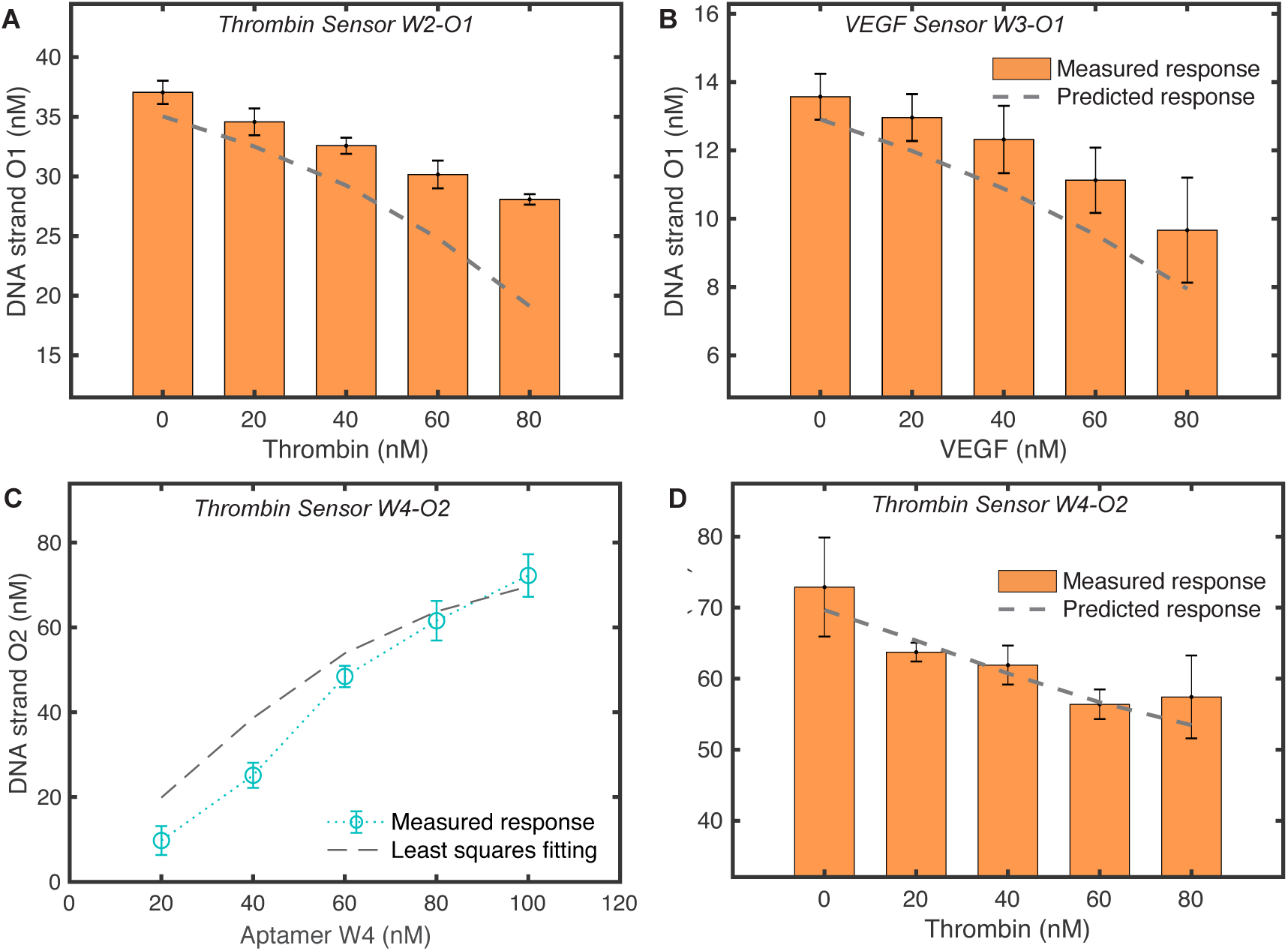
Different exchange process can translate the concentration of different input proteins into the same oligonucleotide output or the same protein into different oligonucleotide outputs. The steady state concentration of the output strand (O1) for **(A)** *Thrombin Sensor W2-O1* in response to different concentrations of thrombin and [W2]=[X2Y2]=[O1Z1] =100 nM; **(B)** *VEGF Sensor W3-O1* in response to different concentrations of VEGF and [W3]= [X3Y3]=[O1Z1] =100 nM. *Thrombin Sensor W2-O1* uses a modified TBA29 DNA aptamer that binds to thrombin (15) while *VEGF Sensor W3-O1* uses a modified 3R02 DNA aptamer that binds to VEGF (35). Even though these exchange processes detect different proteins, they can still produce the same DNA output strand (O1) in response to the input proteins. **(C-D)** *Thrombin Sensor W4-O2* uses the same unmodified DNA aptamer (TBA15) as used *in Thrombin Sensor W1-O1* in the recognition stage, but the translation stage of *Thrombin Sensor W4-O2* uses X4Y4 and O2Z2 complexes which are different then X1Y1 and O1Z1 used in Sensor W1-O1. This enabled us to produce O2 strand from *Sensor W4-O2* that has no sequence relation to the output of *Sensor W1-O1* (SI Table S3 and S8). **(C)** The concentration of O2 produced in response to different concentrations of the W4 strand containing the input aptamer, given 100 nM of each X4Y4 and O2Z2. **(D)** The response of the *Thrombin Sensor W4-O2* to different thrombin input concentrations where [W4] = [X4] = [O2Z2] = 100 nM. As the recognition and the translation stages of *Sensor W4-O2* were modelled similarly as *Thrombin Sensor W1-O1* (see Figure 2 and SI Note S1). For kinetic modelling of the recognition stage for each exchange process (modified TBA29-thrombin and modified 3R02-VEGF), we borrowed the K_d_ value and forward rate constants of original TBA29 (15, 32) and 3R02 aptamers (35). The rates of reactions involved in the translation stage were determined similarly as was for *Thrombin Sensor W1-O1* (see SI Note S1).

### Constructing exchange processes with distinct output signals

So far, the exchange processes we constructed are able to translate different protein concentrations into the same output strand irrespective of aptamers that were used to detect specific proteins. We, therefore, sought to build an alternative exchange process that can produce output strand of different sequences (O2) while incorporating the same protein input and aptamer binding partner (W4) as was in *Thrombin Sensor W1-O1*. This is achieved by modifying TBA15 aptamer with a totally different sequence of d domain compared to earlier designs. In this new exchange process, which is called *Thrombin Sensor W4-O2*, the strand that binds to the thrombin, W4, contains 15-mer TBA15 thrombin aptamer motif and a 15-nucleotide domain at TBA15’s 5’-end of TBA15 that controls which output strand is used. This added domain was, like the domain on W1, designed to minimize interaction with the TBA15 motif that might affect the K_d_ of TBA15-thrombin binding, as in the other sensors, this domain is shared with the output strand sequence. While designing the translation stage for *Thrombin Sensor W4-O2*, we followed the similar design guidelines as we did for *Thrombin Sensor W1-O1* (see Design Section, SI Note S2 and S6). Unlike other sensors, which produces O1 output strand, this exchange process produces O2 strand as an output in Reaction 2.

We followed a similar approach to characterize and predict the response of *Thrombin Sensor W4-O2* as we did for other exchanges process. We first measured the protein free response of *Thrombin Sensor W4-O2* as a function of W4 concentration and then computed k_f,2_ that allows the best fitting between the measured value and model data (Figure 5C). Other rates of the translation stage were calculated similarly as we did for other sensors (SI Note S1). We then measured the response of the exchanges process as a function of thrombin concentration and found the predicted values of O2 are in a close agreement with measured values for different ranges of thrombin concentrations (Figure 5D, SI Fig. S13).

Together, these experiments show a design route to a translation system between protein input concentration and oligonucleotide output concentration where the protein input and oligonucleotide sequence output can be chosen independently. Further, different sensor molecules can be used to detect a particular protein input as long as the aptamer binding is strong enough such that the concentration of free aptamer changes significantly in response to the protein’s presence. We can choose the range of protein input concentrations and oligonucleotide output concentrations by selecting the concentrations of the aptamer and sensor components and use a quantitative model to predict precisely the steady state output concentration of the oligonucleotide that arises in response to a particular protein input concentration.

### Simultaneous detection of different proteins and production of different DNA oligonucleotide outputs

One of the advantages of using DNA aptamers and strand displacement reactions in the exchange process is the structures and sequence-specific interactions between various species of the exchange process. We, therefore, sought to test whether it is possible to configure these exchange process in such a way that they can translate the concertation of different input proteins into a common output signal or the same input protein into different output signals (Figure 1).

We started with testing two sensors with two different protein inputs, and the same oligonucleotide output would respond when operated in the same solution. To do so we studied the responses of the *Thrombin Sensor W2-O1* and *VEGF Sensor W3-O1*, which detect thrombin and VEGF receptively, and both produce the oligonucleotide output O1. For simplicity, we tested how both exchange processes responded to combinations of inputs in a one-pot reaction using both input proteins at the same concentration and varied them by the equal amounts (see Methods). As both of these processes use O1Z1 complex in the translation stage, this limited our ability to track the concentration of the output strands of each process separately when both the of them operated concurrently. We, therefore, relied on the solution of an ODE model that has all the reaction involved in the operation of *Sensor W2-O1* and *W3-O1* concurrently (see SI Note S7).

Because we have shown earlier that the model of each of these sensors can predict their response accurately when operated separately, we would expect to see that the combined model of *Sensor W2-O1* and *W3-O1* should predict the combined response of *Sensor W2-O1* and *W3-O1* as well if *Thrombin Sensor W2-O1* and *VEGF Sensor W3-O1* operate modularly without affecting the operation of the other exchange process. We observed a close match between the measured values and the model data for the combined cases, suggesting that the two-exchange process work almost independently, and we can correctly interpret the concentration of the two input proteins simultaneously and can still predict the output accurately (Figure 6A).

**Figure 6.**
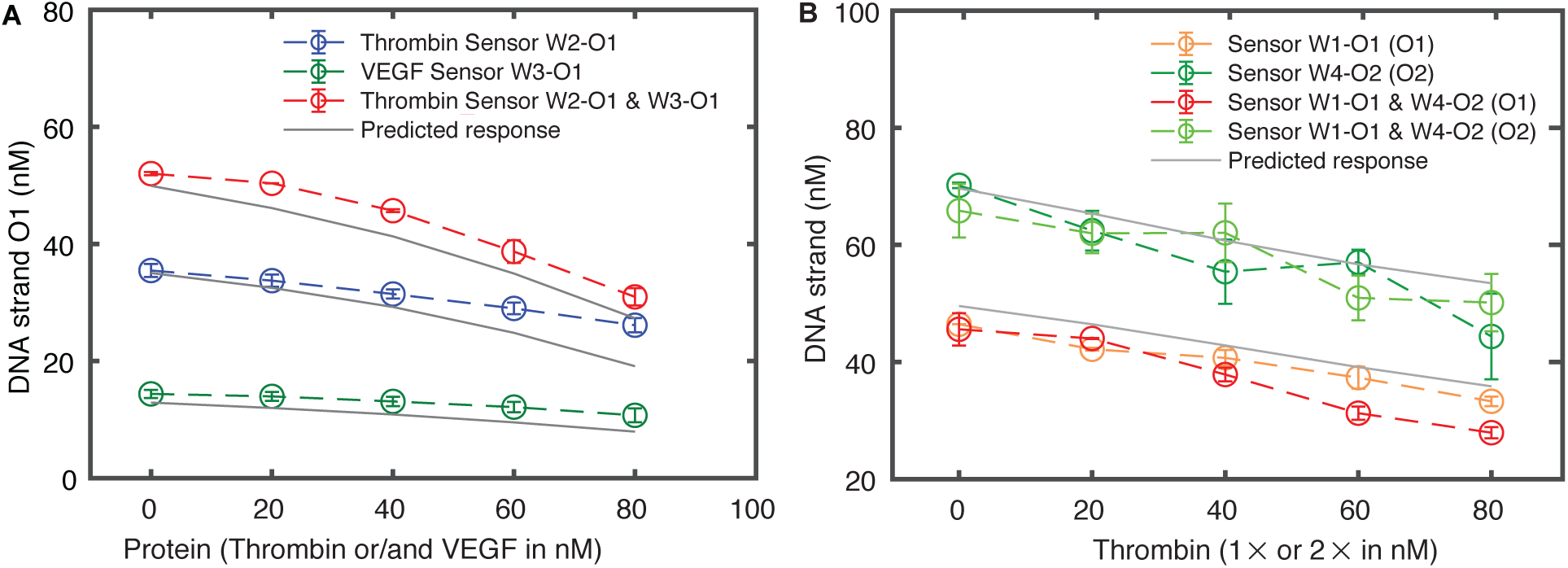
Single output or different outputs can be produced concurrently in response to different proteins or the same protein respectively. *Thrombin Sensor W2-O1 and VEGF Sensor W3-O1* detect thrombin and VEGF respectively, and they produce the same output strand (O1) while *Thrombin Sensor W1-O1* and *Thrombin Sensor W4-O2* both detect thrombin, and they produce different output strands (O1 and O2). **(A)** *Thrombin Sensor W2-O1* and *VEGF Sensor W3-O1* were tested concurrently in the presence [W2]=[W3]=[X2Y2]=[X3Y3]=100 nM and [O1Z1] = 200 nM for different amounts of thrombin and VEGF (0-80 nM) each. As a control individual responses of each exchange process where determined in presence of the respective sensor components ([W2]=[X2Y2]=[O1Z1]=100 nM or [W3]=[X3Y3]= [O1Z1]=100 nM) at different input protein concentrations. The combined response of both the exchange processes was modelled using an ode model that account the same rates that were used to model the individual responses (see SI Note S7 and S1). **(B)** Likewise, the combined response of *Thrombin Sensor W1-O1* (Z1 strand labelled with FAM fluorophore) and *Thrombin Sensor W4-O2* (Z2 strand labelled with HEX fluorophore) was measured in presence of [W1]=[W4]=[X1Y1]=[X4Y4]=[O1Z1]=[O2Z2]=100 nM and compared with the individual responses of each exchange process at 100 nM concentration of the respective sensor components ([W1]=[X1Y1]=[O1Z1]=100 nM or [W4]=[X4Y4]= [O2Z2]=100 nM). In the concurrent operation of Sensor W1-O1 and Sensor W4-O2, thrombin was added two times than the case when the individual response of each exchange process was measured.

We finally sought to examine the operations of the two exchange processes together that can accurately reflect the changes in the same input protein concentration on two DNA output strands that have totally different sequences. For this, we tested the operation of *Thrombin Sensor W1-O1* and *Thrombin Sensor W4-O2* that targets the same protein using the same aptamer but produce, respectively, O1 and O2 strands in response to thrombin protein. Because these processes use totally different OZ complexes in the translation stage, in the combined response, we can track the output of each process separately using different fluorophore molecules attached to the respective Z strand. For this, Z1 and Z2 strands of *Sensor W1-O1* and *W4-O2* were labelled with, respectively, FAM and HEX fluorophore dyes (see Methods).

In this new arrangement, *Thrombin Sensor W4-O2* uses HEX fluorophore dye compared to the previously reported repose of this exchange process that used FAM fluorophore dye. We, therefore, characterized *Thrombin Sensor W4-O2* that used HEX fluorophore dye for different input protein concentrations similarly as was done for *Thrombin Sensor W4-O2* with FAM fluorophore dye. For this, a separate calibration measurement was used to convert the measured fluorescence values into concentrations (see SI Note S3). Subsequently, *Thrombin Sensor W1-O1* and *Thrombin Sensor W4-O2* were tested together in a one-pot reaction in the presence of thrombin that was twice the amount of the cases wherein the response of the individual process was recorded (Figure 7B). We observed that the measured concentration of O1 and O2 strands in one-pot reaction remain close to the same value when individual responses of each sensor were measured separately, suggesting that these two sensors work modularly when operate concurrently.

These results, in general, demonstrate that recognition stage of each of the exchange processes we designed to target only a specific protein, which is governed by the aptamer, and the translation stage of these processes can accurately pass the information of its free aptamer concentration to the output without with minimal crosstalk.

## DISCUSSION

In conclusion, we have constructed a novel modular protein-oligonucleotide exchange process that can produce a DNA oligonucleotide of a specific sequence in response to a protein molecule. The spontaneous interaction of DNA aptamers with specific proteins and their interaction with complementary DNA strands enabled to use fast strand displacement reactions so that an output DNA strand with a limited restriction on it sequence can be produced quickly that reflects the concentration of the input protein. Because the added sequence in the original aptamers should not affect their original operation, there is a limitation on the sequence of DNA output strand. This could be avoided by adding an additional reaction in the translation process such that the sequence of new DNA output strand can be completely independent with the aptamer sequence (SI Fig. S14).

The aptamers we used in this study fold into different confirmations, and that govern their affinity towards the target. As we have modified these aptamers by adding d domain in the original aptamers, this modification might result in increasing or decreasing the affinity of the modified aptamers to the respective proteins. Our mathematical models were able to predict the response of each exchange process accurately. For that, we used the K_d_ values reported in the literature for the unmodified aptamers. This suggests that the modified aptamers can still bind to the respective protein with high affinity. To test this, we varied the K_d_ values for each process in an incremental order while keeping the other rates constant. We found that the modal data for each sensor could not follow the measured response when the K_d_ value was increased by a factor of five (see SI Fig. S15).

Even though our goal was not to construct a sensor that can detect a low concentration of an input protein accurately, we were successfully able to detect 10 nM or higher amounts of different proteins. However, by employing a wide variety of techniques for amplifying the output oligonucleotide, better sensitivity may be achieved (13, 14). Our approach can be easily adapted to sense different protein molecules using their DNA aptamers. To demonstrate this, we designed and characterized four different exchange process that can either target human α-thrombin or VEGF protein molecules. These reaction networks are capable of detecting multiple proteins and producing different DNA output simultaneously.

To model these exchange processes accurately, we proposed a modelling framework wherein the rates of the reactions involved in the translation stage were calculated, and further optimization through experiment or least squares fitting was done to precisely model the aptamer-output response. Using these rates, for each exchange process, we demonstrated that we can accurately predict the output for a given concentration of the protein input. This would allow designing more complex reaction networks that can be used not just to detect different proteins but can also control their activity.

In general, a sensor should not affect the functionality of the sensing molecular. This may be feasible to achieve by realizing different DNA aptamers that bind to a protein at its different binding sites and thereby control different functionality of the protein molecule. For example, two different DNA aptamers of thrombin; TBA15 and TBA29 binds to thrombin but only TBA15 binding to thrombin and alters its enzymatic functionality in the coagulation cascade (33). Therefore, we can use TBA29 for sensing the thrombin concentration and TBA15 to control its functionality.

Each exchange process we designed works well separately and concurrently in a predictable manner, suggesting a robust and reliable operation of these processes and encouraging to adopt this strategy not only to target other proteins but also other biologically relevant molecules such as ATP. Finally, the relative simplicity of our approach inspires to design more complex biochemical systems, which are capable of molecular processing in a sustained manner similar to regulatory pathways.

## MATERIAL AND METHODS

### Storage

All the DNA strands used in this study were ordered from Integrated DNA Technology, Inc. (U.S.A.) in lyophilized form and were PAGE purified except Z1, Z2, O1 and O2 strands which were HPLC purified. O1, O2 and Z1, Z2 strands were labelled with Iowa black FQ quencher and FAM fluorophore dye, respectively, at their 3’ and 5’-ends. For testing the combined operation of *Thrombin Sensor W1-O1* and *Thrombin Sensor W4-O2*, the Z2 strand was labelled with a HEX fluorophore dye. To determine the equilibrium constant of Reaction 2 for *Thrombin Sensor W1-O1*, X1 and Y1 strands were labelled with FAM fluorophore dye and Iowa black FQ quencher at their 5’ and 3’ ends respectively. The lyophilized DNA oligonucleotides were diluted in salt-free water and their concentrations were determined by measuring the absorbance at the wavelength of 260 nm on a standard spectrophotometer using the extinction coefficient provided by IDT for the respective DNA strands. Human **α**-thrombin was purchased from Haematologic Technologies, Inc. (Essex, VT) dissolved in 50% H_2_O/Glycerol. In this paper, we refer to human α-thrombin as thrombin. Recombinant Human VEGF 165 was purchased from R&D Systems Inc. (U.S.A.) in lyophilized form and was reconstituted at 500 μg/mL in sterile 4 mM HCl containing 0.1% bovine serum albumin. Bovine serum albumin lyophilized powder was purchased from Sigma-Aldrich. All the molecules were stored at -20 °C until use.

### Electrophoresis mobility shift assay to study TBA15-thrombin and modified TBA15-thrombin interaction

*Thrombin Sensor W1-O1* uses modified thrombin binding aptamer (TBA) to detect thrombin and in response, modulate the concentration of the output strand O1. To unfold this aptamer into the lowest energy state (27), we prepared 20 µM of modified TBA15 (W1) in 1 × reaction buffer (20 mM Tris, 140 mM NaCl, 1 mM Mg^2+^, 5 mM KCl, and 1 mM CaCl with pH≈7.4) (8, 21) of a total volume 50 µl and annealed it by holding the mixture at 90 °C for 5 min and then gradually cooling down to 25 °C at 1 °C/min rate using Eppendorf Mastercycler. We then added 5 µM of this mixture to separate solutions containing 0, 1, 2.5 and 5 μM thrombin in 1 × reaction buffer components in a total volume of 10 μl. Separate mixtures of original aptamer (TBA15) and thrombin at the same respective concentrations were also made. The samples were incubated for 30 min at 25 °C to allow the binding reaction to approach equilibrium, and then were loaded into a 10% polyacrylamide gel (3.25 ml of 40% polyacrylamide solution, 1.3 ml 10 × reaction buffer, 8.45 ml of MilliQ H_2_O, 60 μl of 10% APS and 6 μl of TEMED). Electrophoresis was performed at 4 °C for 75 min (100 V) in 1 × running buffer (TBE with 5 mM MgCl_2_ and 50 mM KCl). The gels were stained with SYBR Gold (ThermoFisher Scientific) for 30 min and subsequently were imaged using gel imager. To visualize the bands of thrombin, we then stained the gels with EZBlue (Sigma-Aldrich) for 15 min and imaged the gel under white light. Analogous protocols were followed to characterize the reaction of other molecules (b domain of W1Y1 or c domain of X1) with thrombin: pre-annealed W1Y1 complex (5 µM) or X1 strands (5 µM) were incubated without and with thrombin (5 µM) for 30 min at 25 °C and loaded into a 10% polyacrylamide gel. Subsequently, the resulting gel was stained first using SYBR Gold and then with EZBlue.

### Sample preparation for fluorescence spectroscopy experiments

All reactions were performed in 1 × reaction buffer consisting 20 mM Tris, 140 mM NaCl, 1 mM Mg^2+^, 5 mM KCl, and 1 mM CaCl_2_ (with pH≈7.4) unless specified. Different components of the exchange process (W, XY, OZ species) were each pre-annealed at 20 μM concentration in 1 × reaction buffer prior to use by heating the mixes to 90 °C for 5 min and then cooling down to 25 °C at a rate of 1 °C/min, and were kept at 25 °C until use (within a few hours).

### Assay for measuring response in the absence of protein

To measure the aptamer (W)-output (O) response curve for each exchange process, mixtures containing respective 100 nM of each XY nM and OZ pre-annealed complexes (see the aforementioned note) in the reaction buffer were prepared such that the concentration of the pre-annealed respective aptamer (W) will be 20, 40, 60 80 and 100 nM in final solutions of 100 μl for each aptamer concentration. The steady state fluorescence intensities were measured before and after adding the aptamer using FAM filter of Stratagene Mx3000P real-time thermocycler at 25 °C. The changes in the fluorescence intensity that represent the changes in the concentration of the output strand were then converted into concentration using calibration measurements that relate the absolute concentration with the measured fluorescence intensity (see SI Note S3).

### Standard protein sensing assay

To test the response of each exchange process to different concentrations of the input protein (0, 20, 40, 60 and 80 nM), respective input protein (thrombin for Sensor W1-O1, W2-O1, W4-O2, and VEGF for Sensor W3-O1) was incubated with 100 nM of pre-annealed aptamer at 25 °C for 30 min in a total volume of 50 μl containing 1× reaction buffer for each protein concentration. Subsequently, aptamer-protein mixtures were added into 50 μl solutions each containing 100 nM of pre-annealed XY and OZ complexes for that sensor in the reaction buffer for a total volume of 100 µl such that the concentrations of sensor components (W, XY, OZ) were as described in the text. Changes in the fluorescence intensity were recorded using FAM filter of Stratagene Mx3000P real-time thermocycler at 25 °C. The cases wherein the input protein were added in 10 nM and 50 nM incremental changes; the initial concentrations of each sensor component were, respectively, 50 and 500 nM.

### Dynamic protein sensing assay

To test the response of *Thrombin Sensor W1-O1* when thrombin concentration was increased overtime in one test tube, a mixture containing 500 nM of each X1Y1 and O1Z1 complexes in the reaction buffer was prepared and the change in the fluorescence intensity was recorded before and after adding 500 nM of W1 to a total volume of 100 ul. We then added 1ul of a thrombin solution to the mixture so that the mixture contained 100 nM of thrombin and recorded the steady state fluorescence value. The same quantity of thrombin was added several times, and for each addition, steady state fluorescence value was recorded until the final thrombin concentration of the solution reached 400 nM. To test the response of *Thrombin Sensor W1-O1* to decreases in the thrombin concentration overtime in one test tube, we started with 500 nM of X1Y1 and O1Z1 complexes each in reaction buffer and measured the fluorescence intensity and then added 500 nM of W1 and 400 nM of thrombin from the stock solutions in a total volume of 50 μl, and recorded the steady state fluorescence intensity. To decrease the concentration of thrombin without affecting the concentration of the sensor components (W1, X1Y1, O1Z1), 15 μl of a mixture containing 500 nM of each W1, X1Y1, and O1Z1 was added to the solution. The steady state fluorescence value was then recorded. The process of adding W1, X1Y1, and O1Z1 and measuring the steady state fluorescence value was repeated until the final concentration of thrombin reached 182 nM. In both cases, the reported values of the fluorescence were normalized with the fluorescence signal measured when only X1Y1 and O1Z1 complexes were present in the mixtures.

### Comparing dynamic sensing assay with standard sensing assay

To compare the output fluorescence values for dynamic increase and decrease in thrombin concentrations with the standard cases wherein different thrombin concentrations were added in one stage, in the standard cases, we added the same concentration of each molecule as was in the dynamic case for the same total volume but in different aliquots. To compare with dynamic increment cases, 0, 100, 200, 300 and 400 nM of thrombin and 500 nM of W1 (for each case) were added in mixtures containing [X1Y1]= [O1Z1]= 500 nM in a total volume of 100, 101, 102, 103 and 104 ul respectively and recorded the changes in the fluorescence values. To compare with dynamic decrement cases, we added 500 nM of X1Y1 and O1Z1 in aliquots containing 182, 211, 250, 308 and 400 nM of thrombin and W1 of 500 nM (for each case) in a total volume of 120, 95, 80, 65 and 50 ul respectively and the changes fluorescence values were recorded.

### Characterizing the combined response of two exchange networks

To test the combined response of *Thrombin Sensor W2-O1* and *VEGF Sensor W3-O1* to different input proteins, 500 nM of each W2 and W3 aptamers were incubated, respectively, with different amounts of each thrombin and VEGF (0, 100, 200, 300 and 400 nM) at 25 °C for 30 min to a total volume of 50 μl suspended in the reaction buffer. These mixtures were then added into 50 μl solutions containing 500 nM each of X2Y2, X3Y3 and 1000 nM of O1Z1 in the reaction buffer for a total volume of 100 µl such that the concentrations of aptamer and DNA complexes were as described in the text. We then recorded the changes in the fluorescence intensity at 25 °C. A similar protocol was followed for measuring the combined response of *Thrombin Sensor W1-O1* and *Thrombin Sensor W4-O2*. Because Z1 and Z2 strands were labelled with, respectively, FAM and HEX fluorophore dyes, FAM and HEX filters of Stratagene Mx3000P real-time thermocycler were used to measure the changes in fluorescence intensities for *Thrombin Sensor W1-O1 and Thrombin Sensor W4-O2*.

## ACKNOWLEDGEMENT

The authors thank Abdul Majeed Mohammed, Angelo Cangialosi, Dominic Scalise, Josh Fern, and John Zenk for helpful discussions.

## FUNDING

This work was supported in part by the Department of Energy [SC0010595] and the National Science Foundation [CCF-1161941].

