## Supplemental Material for "Modular protein-oligonucleotide signal exchange"

Deepak K. Agrawal

Department of Chemical and Biomolecular Engineering, Johns Hopkins University, 3400 N Charles St, Baltimore, Maryland 21218, USA.

#### Supplementary Note S1: Kinetic modeling of the exchange process

A set of chemical reactions that describe the operation of the exchange process are as follows:

$$W + P \underset{\mathrm{k_{r,1}}}{\overset{\mathrm{k_{f,1}}}{\rightleftharpoons}} WP; \quad K_d = \frac{[W][P]}{[WP]} \tag{S1}$$

$$W + XY \underset{\mathrm{k_{r,2}}}{\overset{\mathrm{k_{f,2}}}{\rightleftharpoons}} WY + X \tag{S2}$$

$$X + OZ \underset{\mathrm{k_{r,3}}}{\overset{\mathrm{k_{f,3}}}{\rightleftharpoons}} XZ + O \tag{S3}$$

Here, aptamer $W$ binds to protein $P$ with a dissociation constant $K_d$. $k_f$ and $k_r$ are the forward and reverse rate constants respectively and the associated subscript with them represent the reaction number. To relate the output concentration with the input concentration accurately, we used mass-action kinetics of Reactions S1-S3 to model the input-output relation at equilibrium:

$$\frac{d[W]}{dt} = k_{r,1}[WP] - k_{f,1}[W][P] + k_{r,2}[X][WY] - k_{f,2}[W][XY], \tag{S4}$$

$$\frac{d[P]}{dt} = k_{r,1}[WP] - k_{f,1}[W][P] \tag{S5}$$

$$\frac{d[WP]}{dt} = -k_{r,1}[WP] + k_{f,1}[W][P], \tag{S6}$$

$$\frac{d[X]}{dt} = k_{f,2}[W][XY] - k_{r,2}[X][WY] + k_{r,3}[O][XZ] - k_{f,3}[X][OZ] \tag{S7}$$

$$\frac{d[XY]}{dt} = -k_{f,2}[W][XY] + k_{r,2}[X][WY] \tag{S8}$$

$$\frac{d[WY]}{dt} = k_{f,2}[W][XY] - k_{r,2}[X][WY] \tag{S9}$$

$$\frac{d[O]}{dt} = -k_{r,3}[O][XZ] + k_{f,3}[X][OZ] \tag{S10}$$

$$\frac{d[OZ]}{dt} = k_{r,3}[O][XZ] - k_{f,3}[X][OZ] \tag{S11}$$

$$\frac{d[XZ]}{dt} = -k_{r,3}[O][XZ] + k_{f,3}[X][OZ] \tag{S12}$$

Here and elsewhere $W$, $P$, $WP$, $XY$, $WY$, $X$, $OZ$, $XZ$ and $O$ species are referred in general for any exchange process, and [] represents the molar concentration of the species. We referred equations S4-S12 as ode model for the exchange process and solved it numerically using Matlab. In this model, $K_d$ and $k_{f,1}$ values of a specific aptamer-protein interaction for an exchange process were taken from the literature (See Supplementary Table S2) and the rate constants of strand displacement reactions ($k_{f,2}$, $k_{r,2}$, $k_{f,3}$, $k_{r,3}$) were calculated using the kinetic rate model of toehold mediated strand displacement reaction, recently proposed by Zhang *et al.* [**?**]. To do that, the kinetic rate model requires accurate calculation of the toehold binding energy for invading and incumbent species involved in a reaction. For example, in Reaction 2, *tb* toehold of *W* strand is invading *XY* complex by displacing the incumbent strand *X* from *XY*. Therefore, to determine the forward rate constant ($k_{f,2}$) of this reaction, we need to calculate the binding energy ($\Delta G$) of the invading strand (*W*) to incumbent complex (*XY*). Similarly, ($k_{f,3}$) requires to calculate the $\Delta G$ of *X* strand binding to *OZ* complex. To calculate the reverse rate constants, for Reactions 2 and 3, ($k_{r,2}$, $k_{r,3}$) *X* and *O* will be the invading strands, and *WY* and *XZ* complexes will be the incumbent complexes respectively. To calculate the $\Delta G$ for each complex, we used NUPACK simulator [**?**] (Supplementary Table S1) and then equations S13- S16 were used to calculated the rates of Reactions 2 and 3 (Supplementary Table S2) [**?**]:

$$\text{Rate Constant} = \frac{k_c k_b k_{\text{incumbent offrate}}}{k_c k_{\text{incumbent offrate}} + k_b k_{\text{invading offrate}} + k_{\text{incumbent offrate}}\, k_{\text{invading offrate}}} \tag{S13}$$

where

$$k_b = \frac{400}{l^2} \tag{S14}$$

$$k_{\text{invading offrate}} = k_c \frac{2}{l} e^{\frac{\Delta G_{\text{invading energy}}}{RT}} \quad \text{(S15)}$$

$$k_{\text{incumbent offrate}} = k_c \frac{2}{l} e^{\frac{\Delta G_{\text{incumbent energy}}}{RT}} \quad \text{(S16)}$$

Here, for Reaction 2, *l* is total number of base-pair of *d* and *c* domains and for Reaction 3, it is the total number of base-pair of *d* domain. $k_c = 3.5 \times 10^6$ $\mathrm{M^{-1}s^{-1}}$, $T = 297$ $K$ and $R$ is the gas constant (8.31 J $\mathrm{mol^{-1}}$ $\mathrm{K^{-1}}$).

Supplementary Table S1: Calculated the binding energy of the complexes involved in the strand displacement reactions. These values were determined using NUPACK simulator [**?**]. A correction factor of 2.38 kcal/mol was added in energies listed here to account the conversion from moles to molar when they were used to calculate the rates [**?**]

| Complex | $\Delta G$ (kcal/mol) *Thrombin Sensor W1-O1* | $\Delta G$ (kcal/mol) *Thrombin Sensor W2-O1* | $\Delta G$ (kcal/mol) *VEGF Sensor W3-O1* | $\Delta G$ (kcal/mol) *Thrombin Sensor W4-O2* |
|---|---|---|---|---|
| *XY* | -25 (*X1Y1*) | –33.42 (*X2Y2*) | -35 (*X3Y3*) | -23.95 (*X4Y4*) |
| *WY* | -25.85 (*W1Y1*) | -34.78 (*W2Y2*) | -35.65 (*W3Y3*) | -24.66 (*W4Y4*) |
| *OZ* | -20.2 (*O1Z1*) | -20.2 (*O1Z1*) | -20.2 (*O1Z1*) | -19.23 (*O2Z2*) |
| *XZ* | -21.7 (*X1Z1*) | -21.64 (*X2Z1*) | -21.32 (*X3Z1*) | -22.14 (*X4Z2*) |

Supplementary Table S2: Rate constants of the strand displacement reactions ($k_{f,2}$, $k_{r,2}$, $k_{f,3}$, $k_{r,3}$) for each exchange network were calculated using Equations S13- S16 [**?**] to model the recognition stage of each exchange process which incorporates aptamer-protein interaction was modelled using $K_d$ and $k_{f,1}$ values that were taken from the literature ($k_{r,1} = K_d k_{f,1}$). The modified values of $k_{f,2}$ (shown inside the brackets) was determined through the least squares fitting of the protein free response of each exchange process. Here, $n$ can be 1, 2, 3 and 4 representing, respectively, *Thrombin Sensor W1-O1*, *Thrombin Sensor W2-O1*, *VEGF Sensor W3-O1* and *Thrombin Sensor W4-O2*.

| Sensor | $K_{d,n}$ (nM) | $k_{f,1,n}$ ($10^4$ $\mathrm{M^{-1}s^{-1}}$) | $k_{f,2,n}$ (modified values) ($10^6$ $\mathrm{M^{-1}s^{-1}}$) | $k_{r,2,n}$ ($10^6$ $\mathrm{M^{-1}s^{-1}}$) | $k_{f,3,n}$ ($10^6$ $\mathrm{M^{-1}s^{-1}}$) | $k_{r,3,n}$ ($10^6$ $\mathrm{M^{-1}s^{-1}}$) |
|---|---|---|---|---|---|---|
| *Thrombin Sensor W1-O1* | 5.2[?] | 7.45[?] | 2.82 (0.065) | 0.67 | 3.24 | 0.26 |
| *Thrombin Sensor W2-O1* | 0.5[?] | 2[?] | 3.18 (0.005) | 0.32 | 3.22 | 0.28 |
| *VEGF Sensor W3-O1* | 0.3[?] | 1.92[?] | 2.62 (0.0004) | 0.88 | 3.04 | 0.46 |
| *Thrombin Sensor W4-O2* | 5.2[?] | 7.45[?] | 2.69 (0.08) | 0.81 | 3.47 | 0.03 |

### Supplementary Note S2: Designing Thrombin Sensor W1-O1

Reliable operation of the exchange process requires that there is minimal crosstalk between species that are not programmed to interact. In particular, when the aptamer forms *W1Y1* complex by displacing *X1* from *X1Y1* in Reaction S2, the aptamer should no longer be able to bind to the input protein. Specifically, this means that the *W1Y1* complex should not interact with the protein through the *b* domain of the aptamer. Such an interaction could lower the concentration of the output strand (See Figure 2 in the main paper). Likewise, the *X1* strand that is released from this reaction contains *c* domain which is presented in single-stranded form. It is also critical that this *X1* strand not be able to interact with the input protein. These interactions may be avoided by choosing short *b* and *c* domains.

Further, as we require that any change in concentration of the *W1* strand should quickly be reflected in the output, the length of toehold domains should be four base-pair or longer in order to allow Reactions S2 and S3 to reach the equilibrium quickly. For reversible toehold-mediated strand-displacement reactions, the rate constants can typically vary from 1 $M^{-1}s^{-1}$ (zero toehold ) to $6 \times 10^6$ $M^{-1}s^{-1}$ (7 toehold or longer) [**?**]. Finally, *tc* domain should not share any complementary regions with *c* domain of the aptamer in order to prevent the reversibility of Reaction 3. Based on these and other design considerations (See Design Section in the main paper), we designed *Thrombin Sensor W1-O1* to exchange thrombin concentration into output strand *O1* using a modified TBA15 DNA aptamer [**?**].

Supplementary Table S3: Sequences of the DNA strands used in *Thrombin Sensor W1-O1* that uses modified TBA15 DNA aptamer to sense input protein thrombin. Here, domains *c*, *tb* and *b* represent the sequence of the original aptamer which folds into a two stacked G-quadruplexes and binds to thrombin with a dissociation constant ($K_d$) of 5.2 nM [**?**, **?**].

| Name of the DNA Strand | Domains | Sequence |
|---|---|---|
| *W1* | d c tb b | TAATTTATAATTATT GG TTGGT GTGGTTGG |
| *X1* | td d c | ATTATG TAATTTATAATTATT GG |
| *Y1* | tb* c* d* td* | ACCAA CC AATAATTATAAATTA CATAAT |
| *Z1* | F tc* d* td* | /56-FAM/ CTCTT AATAATTATAAATTA CATAAT |
| *O1* | d tc Q | TAATTTATAATTATT AAGAG /3IABkFQ/ |

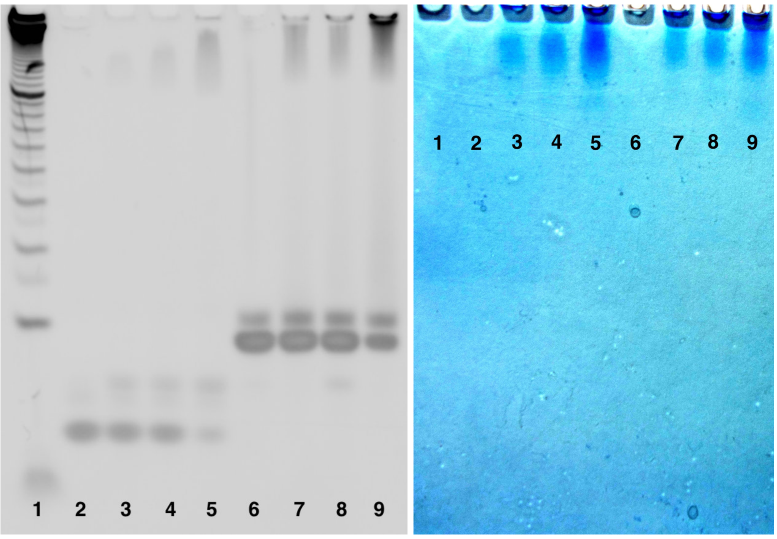


(A) (B)

Supplementary Figure S1: **Electrophoresis mobility shift assay for characterizing the interaction of thrombin with the modified TBA15 DNA aptamer**. Lane 1 is a 10 base-pair ladder (ThermoFisher Scientific) while lane 2-5 and 6-9, respectively, indicating a fixed concentration of pre-annealed 15-mer TBA15 and modified TBA15 (*W1*) aptamers that were incubated with different amounts of thrombin (See Supplemental Table S4) at 25 $^{o}$C for 30 min before loading the samples into the gel (See Method Section in the main paper) (A) Detection of the DNA with SYBR Gold staining. (B) Detection of thrombin with EZBlue staining. An increase in thrombin concentration increases its migration in the presence of 15-mer TBA15 or modified TBA15 aptamer by almost the same amount, suggesting the affinity of the modified aptamer towards thrombin may not be significantly altered compared to the original aptamer.

Supplementary Table S4: Stoichometry of the gel matrix shown in Supplemental Figure S1.

| Lane | Original TBA15 asptamer ($\mu$M) | Thrombin protein ($\mu$M) | Modified TBA15 aptamer ($\mu$M) |
|---|---|---|---|
| 2 | 5 | 0 | - |
| 3 | 5 | 1 | - |
| 4 | 5 | 2.5 | - |
| 5 | 5 | 5 | - |
| 6 | - | 0 | 5 |
| 7 | - | 1 | 5 |
| 8 | - | 2.5 | 5 |
| 9 | - | 5 | 5 |

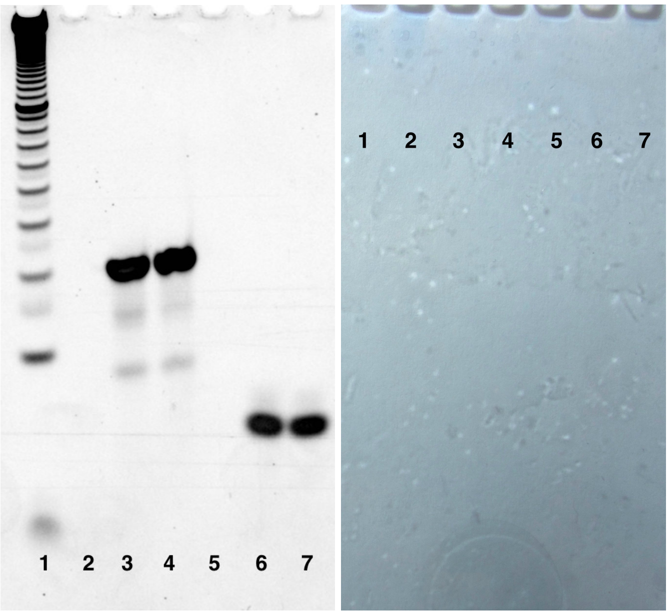


(A) (B)

Supplementary Figure S2: **Electrophoresis mobility shift assay to detect the interaction between *b* or *c* domain with thrombin**. Lane 1 is a 10 base-pair ladder (ThermoFisher Scientific), lane 2-4 indicating different concentrations of thrombin and *W1Y1* complex, which has a single stranded *b* domain (See Figure 2 in the main text), in the ratio of 1:0, 0:1 and 1:1 while lane 5-7 indicating thrombin and *X1* strand, which has *c* domain, in the ratio of 1:0, 0:1 and 1:1 respectively (See Supplemental Table S5) (A) Detection of the DNA with SYBR Gold staining. (B) Detection of thrombin with EZBlue staining. In the gel matrix, no bands of thrombin were observed in the presence of *W1Y1* complex or *X1* strand, suggesting that *b* or *c* domain alone has no affinity towards thrombin.

Supplementary Table S5: Stoichometry of the gel matrix shown in Supplemental Figure S2.

| Lane | Protein ($\mu$M) | *W1Y1* ($\mu$M) | Thrombin ($\mu$M) | *X1* ($\mu$M) |
|---|---|---|---|---|
| 2 | 5 | 0 | - | - |
| 3 | 0 | 5 | - | - |
| 4 | 5 | 5 | - | - |
| 5 | - | - | 5 | 0 |
| 6 | - | - | 0 | 5 |
| 7 | - | - | 5 | 5 |

#### Supplementary Note S3: Calibration fluorophore dyes to determine the concentration of the output strand from the measured responses

In this study, while measuring the responses of each exchange process without or with protein, we recorded the changes in the fluorescence intensity before and after adding the input molecule (aptamer or protein respectively). We then normalized the measured fluorescence signal with respect to background which was measured in the absence of the input molecule. The normalized value corresponds to a specific concentration of *XZ* complex present in the reaction volume at the equilibrium. In order to compare the experimental data with the model data accurately, we calibrated the fluorophore molecule so that we can relate the measured fluorescence signal with the exact concentration of *XZ* complex.

To determine the relation between fluorescence signal and [*XZ* ], we need to know the exact amount of *XZ* present in a reaction mixture at the equilibrium. Because in the current design of translation stage, Reaction 3 is reversible, it is not straightforward to relate the output [*XZ*] with input [*X*] at the equilibrium without knowing the exact rate constants for these reaction. Therefore, to reduce the number of unknown parameters, we used a DNA strand (*Z1**) that can form a duplex with *Z1* by displacing *O1* from *O1Z1* complex (Supplemental Figure S3). Because *Z1** is completely complementary to *Z1*, this reaction is irreversible as no toehold will be available for *O1* to interact with *Z1***Z1* duplex, once *Z1** strand hybridized with *Z1*. At the equilibrium, the concentration of *Z1***Z1* complex must be equal to the input concentration of *Z1**. Therefore, by diving the measured normalized fluorescence signal with the input concentration of *Z1**, we can calculate the average calibration factor that relates the concentration of *Z1***Z1* with the measured fluorescence signal.

To determine the calibration factor, a fixed concentration of pre-annealed *O1Z1* complex (500 nM) was mixed with different concentrations *Z1** (50, 100, 150, 200 and 250 nM) and the changes in the fluorescence intensity were recorded before and after adding *Z1** (Supplemental Figure S4A) and average calibration factor was calculated. Similarly experiments were performed to calibrate the FAM and HEX fluorophore labeled complex of *Thrombin Sensor W4-O2* (Supplemental Figure S4B and S5) to determine the calibration factor. Using these values, we converted the concentration of the input *Z1** strand in fluorescence unit and a close agreement between the measured and the calculated responses were observed (Supplemental Figure S4 and S5).

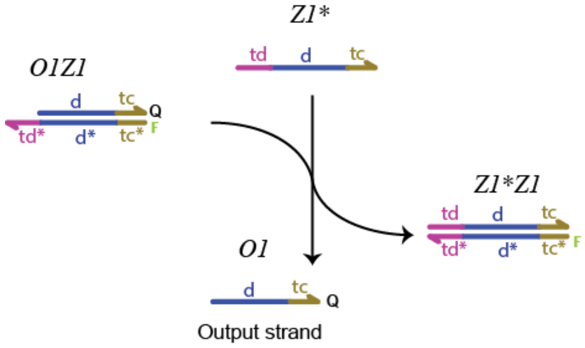


Supplementary Figure S3: Irreversible strand displacement reaction to characterize the measured fluorescence intensity and the concentration of the presented fluorophore labeled complex.

Due to the fact that any change in the [*XZ*] would also represent the same change in the [*O*] strand, in this work, we reported the measured response as change in the concentration of the output strand. Further, for each experiment, we ran similar calibration measurements as mentioned above so that the measured fluorescence signal can be converted into concentration accurtaly.

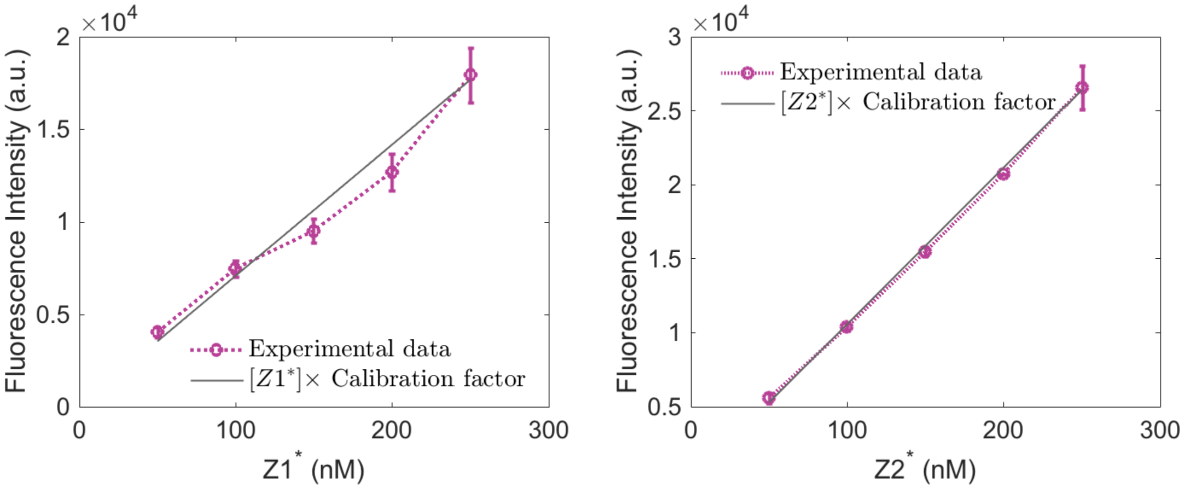


(A) (B)

Supplementary Figure S4: **Fluorescence calibration curves of FAM fluorophore.** Variations in the fluorescence intensity were recorded before and after adding different amounts of respective $Z^*$ strand in the mixtures where the respective [*OZ*] =500 nM (*Z1** and *O1Z1* for (A), and *Z2** and *O2Z2* for (B)). (A) For *Thrombin Sensor W1-O1*, *Thrombin Sensor W2-O1* and *VEGF Sensor W3-O1*, a fluorescence unit is equal to 14.26 pM of $Z^*$ strand while (B) for *Thrombin Sensor W4-O2*, it is 9.47 pM. It should be noted that *Thrombin Sensor W1-O1*, *Thrombin Sensor W2-O1* and *VEGF Sensor W3-O1* use the same *O1Z1* complex compared to *Thrombin Sensor W4-O2* that uses *O2Z2* complex that is totally different then *O1Z1*.

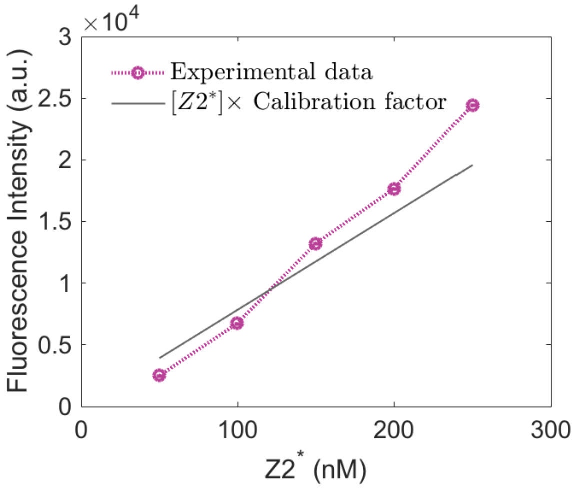


Supplementary Figure S5: **Fluorescence calibration curve of HEX fluorophore used in Thrombin Sensor W4-O2.** Variations in the fluorescence intensity were recorded before and after adding different amounts of *Z2** strand in the mixtures having [*O2Z2*]=500 nM. For HEX fluorophore, a fluorescence unit is equal to 13.54 pM of *Z2** strand.

### Supplementary Note S4: Measuring the equilibrium constant of Reaction 2 for Thrombin Sensor W1-O1

For the calculated value of $k_{f,2}$, we found that ode model of the exchange process is not able predict accurately the measured response of *Thrombin Sensor W1-O1* in absence of thrombin (Supplementary Figure S6). This may be due to the G-quadruplex folding of the TBA15 aptamer that may result in reducing its interaction with the *X1Y1* complex through *tb* toehold. To test this hypothesis, we used the modified *X1Y1* complex of *Sensor W1-O1* by labelling the *X1* and *Y1* strands with a fluorophore and a quencher molecule respectively (Supplementary Figure 7A, See Methods) and then measured the changes in the fluorescence intensity with different amounts of *W1* strand (Supplementary Figure 7B). We found a best fit between the measured response and model data for a $k_{f,2}$ value of 5.02 $\times 10^4$ $M^{-1}s^{-1}$ which is much smaller than the calculated value (2.82 $\times 10^6$ $M^{-1}s^{-1}$).

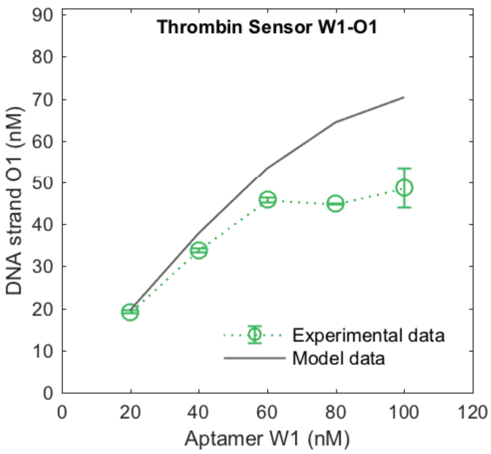


Supplementary Figure S6: . Comparison of measured protein free response with the model data when the calculated value of $k_{f,2}$ (2.82 $\times$ $10^6$ $M^{-1}s^{-1}$) was used (See Supplementary Table S2).

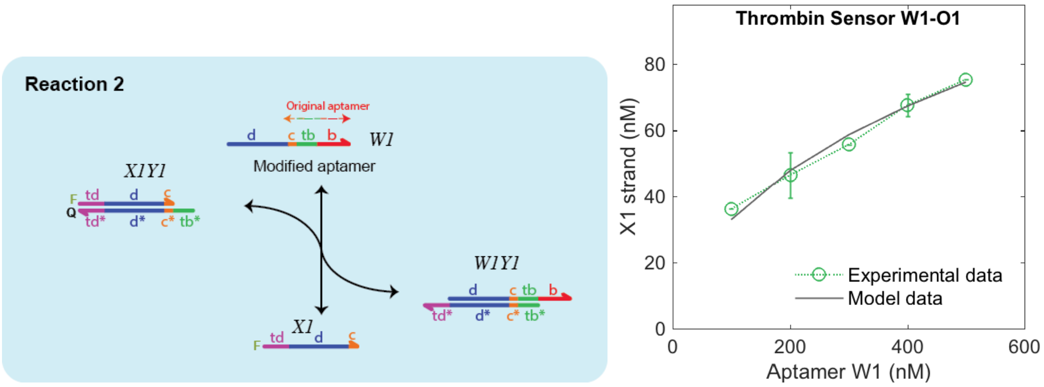


(A) (B)

Supplementary Figure S7: **Determining the forward rate constant of Reaction 2.** (A) A schematic representation of Reaction 2 alone where *X1Y1* complex was labelled with a fluorophore dye and a quencher (See Methods) so that any change in the [*X1*] can be directly detected when *W1* displaces *X1* from *X1Y1* complex. (B) We added different concentrations of *W1* in mixtures that have 250 nM of *X1Y1* and changes in the fluorescence intensity were recorded before and after adding the *W1* strand. These changes were then converted into concentration through directly calibrating the fluorescence to [*X1*] relation. A best fit between the measured and the model data was found when $k_{f,2}$ = 5.02 $\times 10^4$ $M^{-1}s^{-1}$.

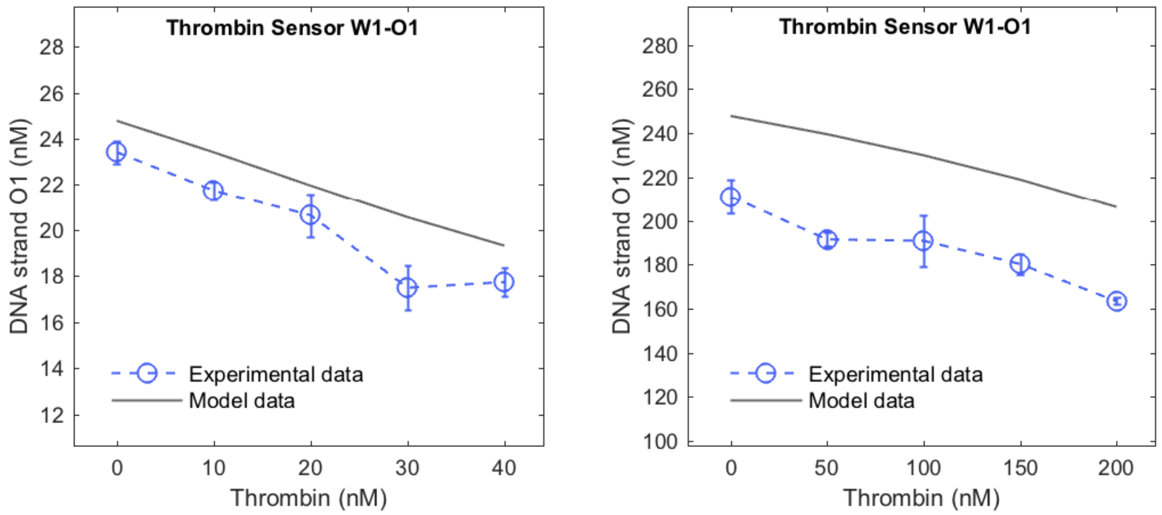


(A) (B)

Supplementary Figure S8: **Predictable response of Thrombin Sensor W1-O1 at different rages of input protein concentrations**. Measured response of *Thrombin Sensor W1-O1* where thrombin was added in different incremental amounts in the presence of (A) [*W1*]=[*X1Y1*]=[*O1Z1*]=50 nM, and (B) [*W1*]=[*X1Y1*]=[*O1Z1*]=500 nM. For each variation in thrombin concentration, model data can track the output.

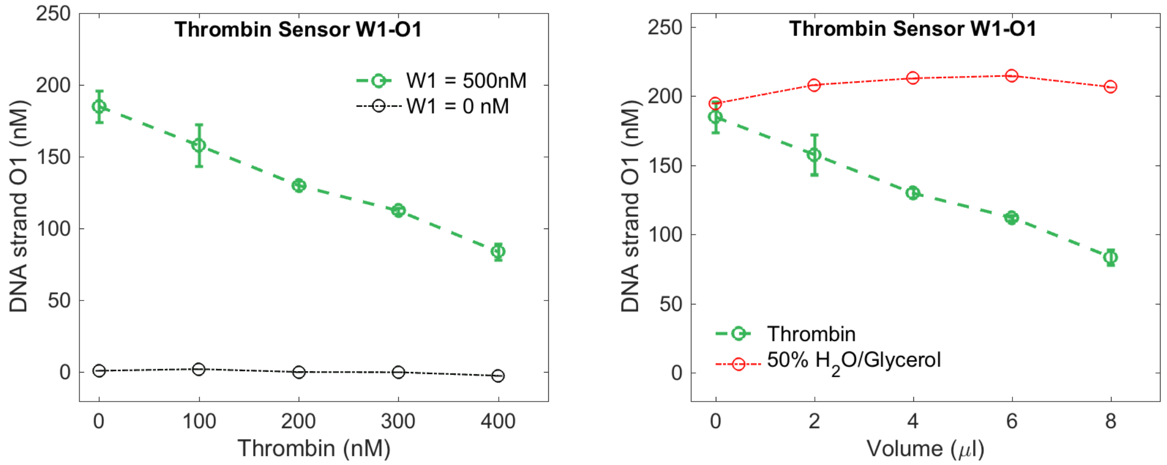


(A) (B)

Supplementary Figure S9: **Thrombin Sensor W1-O1 operates reliably**. (A) Measured response of *Thrombin Sensor W1-O1* in the presence and absence of *W1* strand (500 nM) and [*X1Y1*]=[*O1Z1*]=500 nM. In the absence of *W1*, concentration of the output strand does not change significantly with different amounts of thrombin, suggesting that *X1Y1* or *O1Z1* do not interact with the thrombin. (B) When 50% $H_2O$/Glycerol mixture was added instead of thrombin in the presence of [*W1*]=[*X1Y1*]= [*O1Z1*]=500 nM, no significant change in the output signal was observed with respect to the value when no 50% H2O/Glycerol was added, suggesting that $H_2O$/Glycerol base mixture in which thrombin was dissolved does not affect the operation of the exchange process adversely.

#### Supplementary Note S5: Designing Thrombin Sensor W2-O1 and VEGF Sensor W3-O1

For *Thrombin Sensor W2-O1* and *VEGF Sensor W3-O1*, we added the same 15 nucleotide sequence (*d* domain) at the 5'-ends of TBA29 and 3R02 aptamers as used for *Thrombin Sensor W1-O1*. This enabled us to produce the same *O1* strand for *Thrombin Sensor W2-O1* and *VEGF Sensor W3-O1* as in *Thrombin Sensor W1-O1*, even though, they use totally different aptamers and/or sense different proteins. We then group the core sequence of each aptamer into *c*, *tb* and *b* domains while minimizing the possibility of *WY* or *X* species binding with the input protein through *b* or *c* domain respectively for each exchange process. This is achieved by allowing limited base-pairs for overhang *b* domain and a short *c* domain. We use the same sequence of the toehold *td* for each *Y* strand for *Thrombin Sensor W2-O1* and *VEGF Sensor W3-O1* as used for *Thrombin Sensor W1-O1*, so that the same *O1Z1* complex could be used for *Thrombin Sensor W1-O1*, *Thrombin Sensor W2-O1* and *VEGF Sensor W3-O1*.

Supplementary Table S6: Sequences of the DNA strands used in *Thrombin Sensor W2-O1* which uses modified TBA29 DNA aptamer to target thrombin. Here, domains *c*, *tb* and *b* represent the sequence of the original aptamer that binds to thrombin with a dissociation constant ($K_d$) of 0.5 nM when the confirmation change occurs [?].

| Name of the DNA Strand | Domains | Sequence |
|---|---|---|
| W2 | d c tb b | TAATTTATAATTATT AGTCCGTG GTAGG GCAGGTTGGGGTGACT |
| X2 | td d c | ATTATG TAATTTATAATTATT AGTCCGTG |
| Y2 | tb* c* d* td* | CCTAC CACGGACT AATAATTATAAATTA CATAAT |
| Z1 | F tc* d* td* | /56-FAM/ CTCTT AATAATTATAAATTA CATAAT |
| O1 | d tc Q | TAATTTATAATTATT AAGAG /3IABkFQ/ |

Supplementary Table S7: Sequences of the DNA strands used in *VEGF Sensor W3-O1* which uses modified 3R02 DNA aptamer to target VEGF. Here, domains *c*, *tb* and *b* represent the sequence of the original aptamer that binds to VEGF with a dissociation constant ($K_d$) of 0.3 nM when the confirmation change occurs [?].

| Name of the DNA Strand | Domains | Sequence |
|---|---|---|
| W3 | d c tb b | TAATTTATAATTATT TGTGGGGG TGGA CTGGGTGGGTACC |
| X3 | td d c | ATTATG TAATTTATAATTATT TGTGGGGG |
| Y3 | tb* c* d* td* | TCCA CCCCCACA AATAATTATAAATTA CATAAT |
| Z1 | F tc* d* td* | /56-FAM/ CTCTT AATAATTATAAATTA CATAAT |
| O1 | d tc Q | TAATTTATAATTATT AAGAG /3IABkFQ/ |

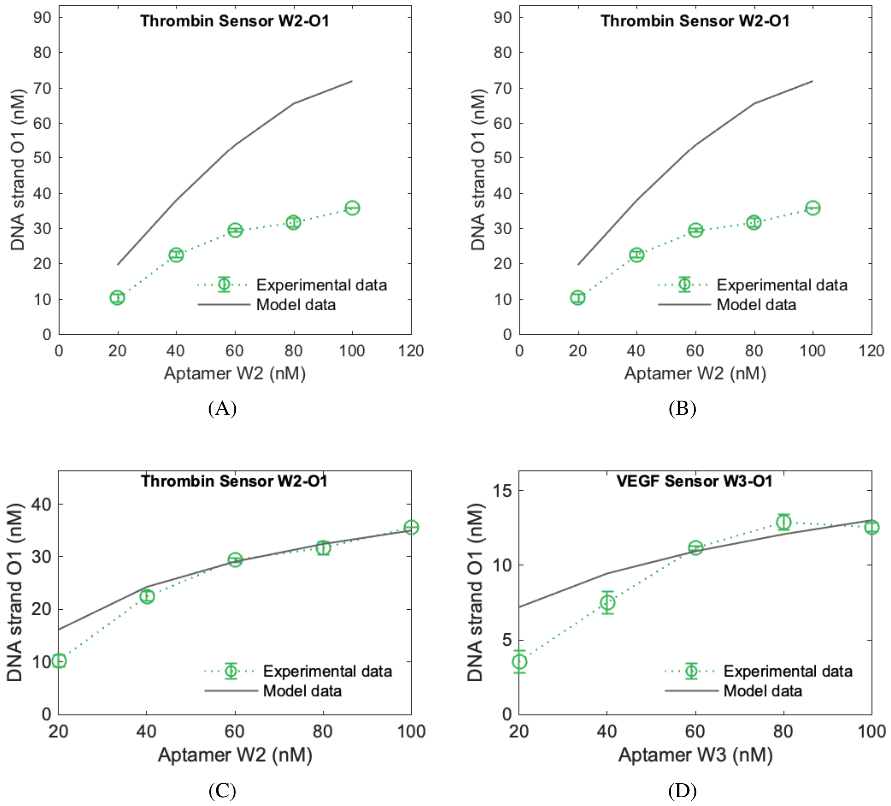


Supplementary Figure S10: **Thrombin Sensor W2-O1 and VEGF Sensor W3-O1 can relate the free aptamer concentration with the output signal.** *Thrombin Sensor W2-O1* uses the modified TBA29 aptamer (*W2*) as a thrombin binding aptamer and thrombin as a target protein while *VEGF Sensor W3-O1* uses modified 3R02 aptamer (*W3*) to detect VEGF as a target protein. Protein free response of (A, C) *Sensor W2-O1* where [*X2Y2*]= [*O1Z1*]=100 nM and (B, D) *Sensor W3-O1* when [*X3Y3*]= [*O1Z1*]=100 nM. In each case, ode model of the exchange process was used to predict these response while using the $k_{f,2}$ that was either (A, B) calculated or (C, D) found through the least squares fitting (See Supplementary Note S1 for details).

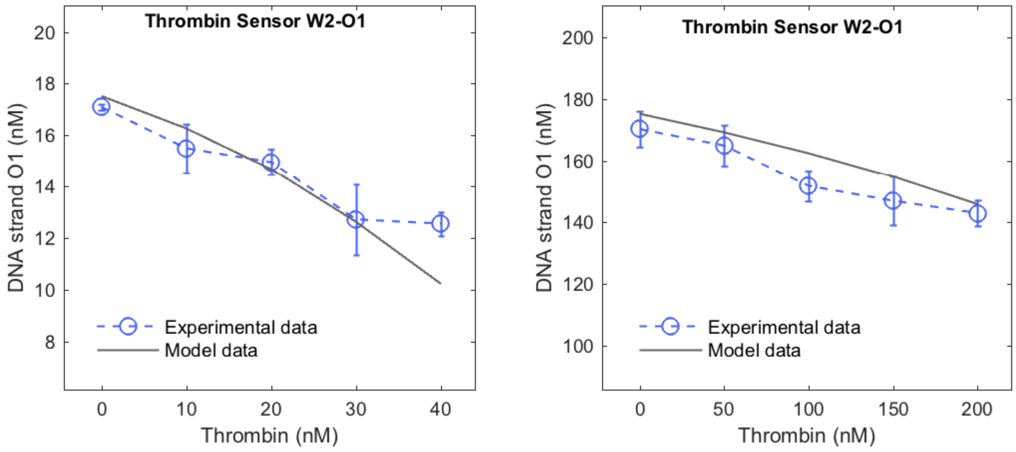


(A) (B)

Supplementary Figure S11: **A reliable predictable response of Thrombin Sensor W2-O1 at different rages of input protein concentrations**. Measured response of *Thrombin Sensor W2-O1* where thrombin was added in different incremental amounts in the presence of (A) [*W2*]=[*X2Y2*]=[*O1Z1*]=50 nM, and (B) [*W2*]=[*X2Y2*]=[*O1Z1*]=500 nM. For each variation in thrombin concentration, model data can track the output.

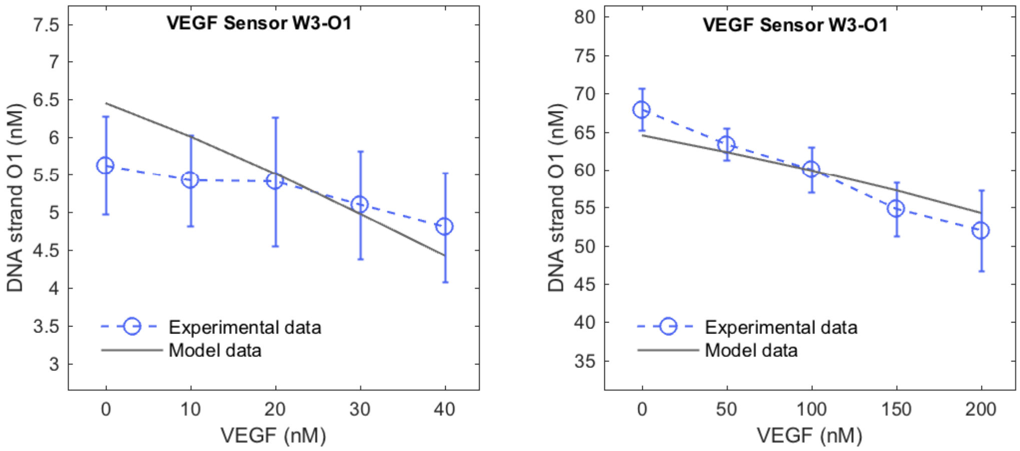


(A) (B)

Supplementary Figure S12: **A reliable predictable response of VEGF Sensor W3-O1 at different rages of input protein concentrations**. Measured response of *VEGF Sensor W3-O1* where thrombin were added in different incremental amounts in the presence of (A) [*W3*]=[*X3Y3*]=[*O1Z1*]=50 nM, and (B) [*W3*]=[*X3Y3*]=[*O1Z1*]=500 nM. For each variation in VEGF concentration, model data can track the output.

#### Supplementary Note S6: Designing Thrombin Sensor W4-O2

*Thrombin Sensor W4-O2* take thrombin as in input protein and passes this information to an output strand *O2* that has no sequence in common with the output strand *O1* of other exchange processes we designed so far. This is achieved by changing the sequence of *d* and *tc* domains compared to *Thrombin Sensor W1-O1*. They both uses the same sequence of the original TBA15 aptamer (See Table 1 in the main text).

Supplementary Table S8: Sequences of the DNA strands used in *Thrombin Sensor W4-O2* that uses TBA15 DNA aptamer to target thrombin but produces a different output strand (*O2*) compared to output strand (*O1*) of other three sensors.

| Name of the DNA Strand | Domains | Sequence |
|---|---|---|
| *W4* | d c tb b | TAATATATAATAATA GG TTGGT GTGGTTGG |
| *X4* | td d c | TTAATG TAATATATAATAATA GG |
| *Y4* | tb* c* d* td* | ACCAA CC TATTATTATATATTA CATTAA |
| *Z2* | F tc* d* td* | /56-FAM/ TTTTC TATTATTATATATTA CATTAA |
| *O2* | d tc Q | TAATATATAATAATA GAAAA /3IABkFQ/ |

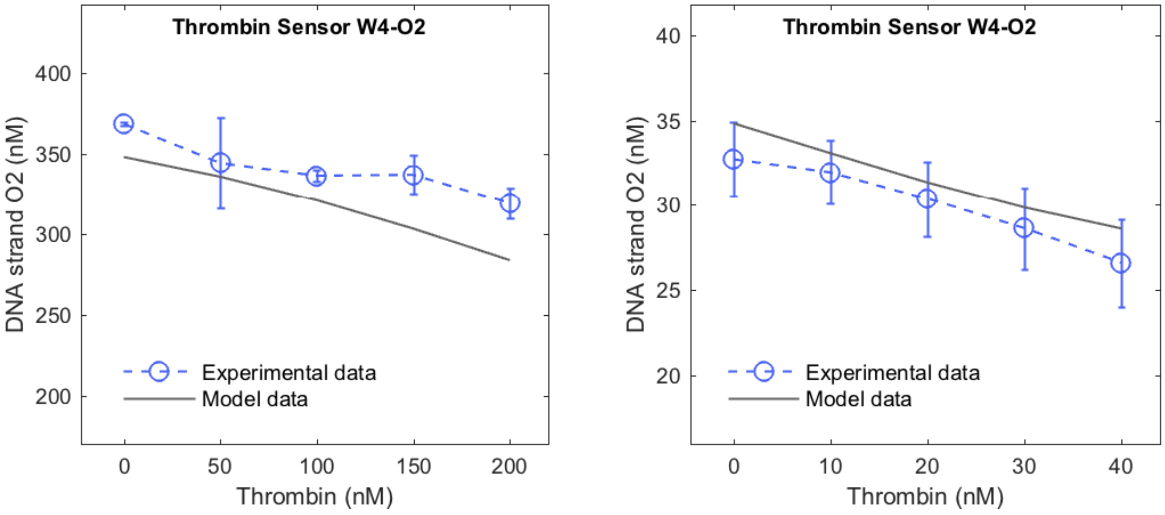


(A) (B)

Supplementary Figure S13: **A reliable predictable response of Thrombin Sensor W4-O2 at different rages of input protein concentrations**. Measured response of *Thrombin Sensor W4-O2* where thrombin were added in different incremental amounts in the presence of (A) [*W4*]=[*X4Y4*]=[*O2Z2*]=50 nM, and (B) [*W2*]=[*X4Y4*]=[*O2Z2*]=500 nM. For each variation in thrombin concentration, model data can track the output.

#### Supplementary Note S7: Modeling combined response of Thrombin Sensor W2-O1 and VEGF Sensor W3-O1

To model the combined response of *Thrombin Sensor W2-O1* and *VEGF Sensor W3-O1*, we used the following set of reactions:

$$\mathrm{W2} + \mathrm{P2} \underset{\mathrm{k_{r,1,2}}}{\overset{\mathrm{k_{f,1,2}}}{\rightleftharpoons}} \mathrm{W2\,P2};\ K_{d,2} = \frac{[W2][P2]}{[W2P2]} \tag{S17}$$

$$\mathrm{W2} + \mathrm{X2\,Y2} \underset{\mathrm{k_{r,2,2}}}{\overset{\mathrm{k_{f,2,2}}}{\rightleftharpoons}} \mathrm{W2\,Y2} + \mathrm{X2} \tag{S18}$$

$$\mathrm{X2} + \mathrm{O1\,Z1} \underset{\mathrm{k_{r,3,2}}}{\overset{\mathrm{k_{f,3,2}}}{\rightleftharpoons}} \mathrm{X2\,Z1} + \mathrm{O1} \tag{S19}$$

$$\mathrm{W3} + \mathrm{P3} \underset{\mathrm{k_{r,13,}}}{\overset{\mathrm{k_{f,1,3}}}{\rightleftharpoons}} \mathrm{W3\,P3};\ K_{d,3} = \frac{[W3][P3]}{[W3P3]} \tag{S20}$$

$$\mathrm{W3} + \mathrm{X3\,Y3} \underset{\mathrm{k_{r,2,3}}}{\overset{\mathrm{k_{f,2,3}}}{\rightleftharpoons}} \mathrm{W3\,Y3} + \mathrm{X3} \tag{S21}$$

$$\mathrm{X3} + \mathrm{O1\,Z1} \underset{\mathrm{k_{r,3,3}}}{\overset{\mathrm{k_{f,3,3}}}{\rightleftharpoons}} \mathrm{X3\,Z1} + \mathrm{O1} \tag{S22}$$

Here, $k_{f1,2}, k_{r,1,2}, k_{f2,2}, k_{r,2,2}, k_{f3,2}, k_{r,3,2}$ and $k_{f1,3}, k_{r,1,3}, k_{f2,3}, k_{r,2,3}, k_{f3,3}, k_{r,3,3}$ are the rates for *Thrombin Sensor W2-O1* and *VEGF Sensor W3-O1* respectively. Using mass-action kinetics, we derived the coupled differential equations model of Reaction S17-S22 (as was done in Supplementary Note S1) and solved them numerically while using the rates shown in Supplementary Table S2.

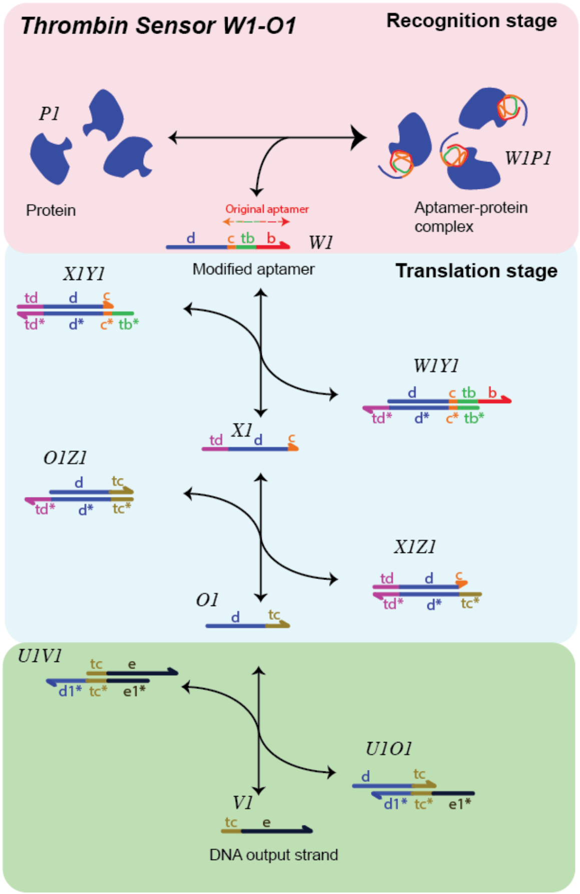


Supplementary Figure S14: **Modified exchange process to produce a DNA output strand of completely independent sequence from the original aptamer's sequence** . By adding an additional strand displacement reaction (green) into the existing translation stage (cyan), a new DNA output strand (*V1*) can have any sequence as the sequence of the added the *d* domain which is added in the original aptamers, will not be a part of the *V1* strand. *V1* would still be able to reflect the concentration of the input protein.

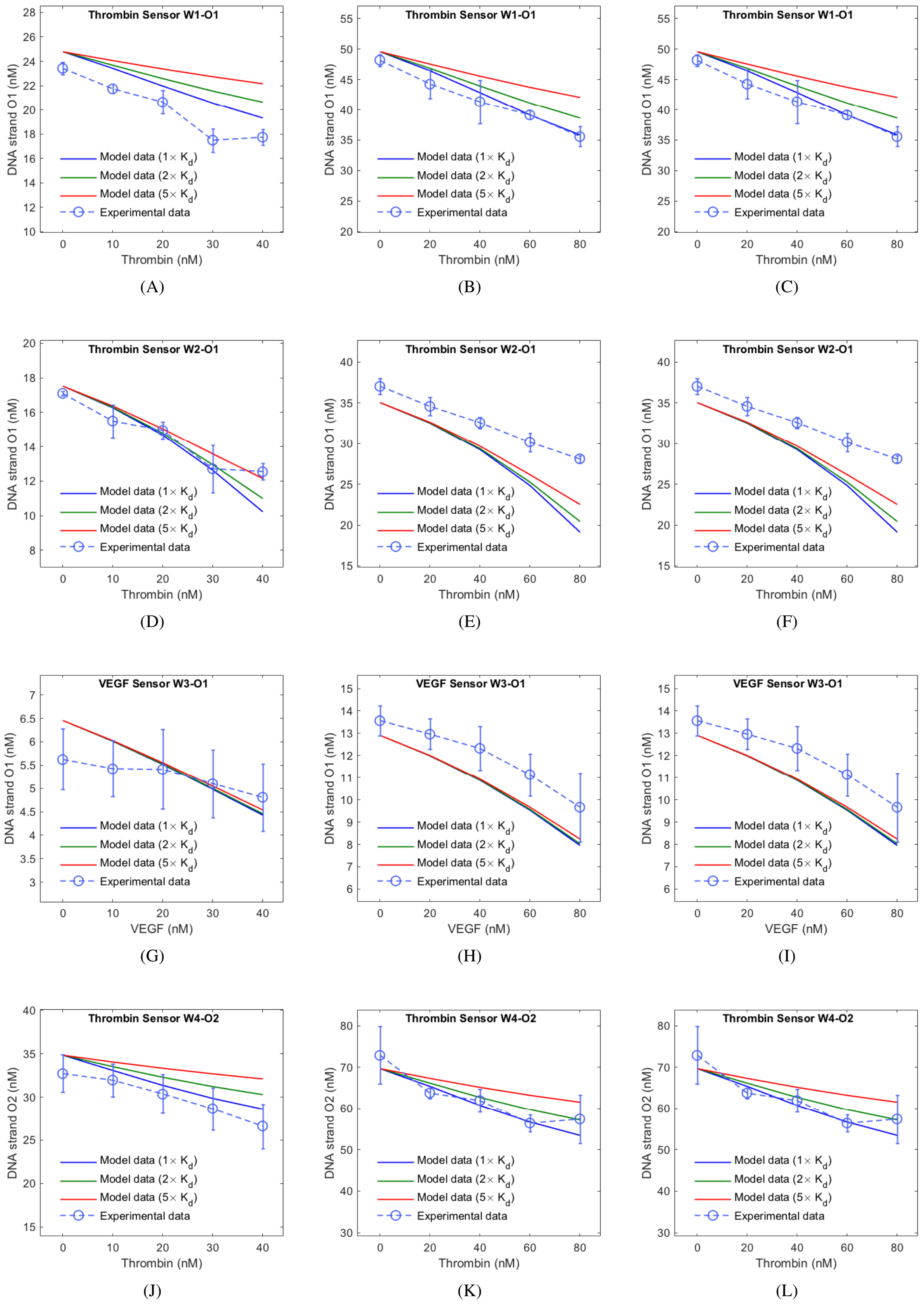


Supplementary Figure S15: **Variation in $K_d$ values does not significantly effect the model data**. Affect of different $K_d$ values on the model data for (A-C) *Thrombin Sensor W1-O1*, (D-F) *Thrombin Sensor W2-O1*, (G-I) *VEGF Sensor W3-O1* and (J-L) *Thrombin Sensor W4-O2* are compared with the measured responses.
